# Downscaling mutualistic networks from species to individuals reveals consistent interaction niches and roles within plant populations

**DOI:** 10.1101/2024.02.02.578595

**Authors:** Elena Quintero, Blanca Arroyo-Correa, Jorge Isla, Francisco Rodríguez-Sánchez, Pedro Jordano

## Abstract

Species-level networks emerge as the combination of interactions spanning multiple individuals, and their study has received considerable attention over the past 30 years. However, less is known about the structure of interaction configurations within species, even though individuals are the actual interacting units in nature.

We compiled 46 empirical, individual-based, interaction networks on plant-animal seed dispersal mutualisms, comprising 1037 plant individuals across 29 species from various regions. We compared the structure of individual-based networks to that of species-based networks and, by extending the niche concept to interaction assemblages, we explored individual plant specialization. Using a Bayesian framework to account for uncertainty derived from sampling, we examined how plant individuals “explore” the interaction niche of their populations.

Both individual-based and species-based networks exhibited high variability in network properties, lacking remarkable structural and topological differences between them. Within populations, frugivores’ interaction allocation among plant individuals was highly heterogeneous, with one to three frugivore species dominating interactions. Regardless of species or bioregion, plant individuals displayed a variety of interaction profiles across populations, with a consistently small percentage of individuals playing a central role and exhibiting high diversity in their interaction assemblage. Plant populations showed variable mid to low levels of niche specialization; and individuals’ interaction niche “breadth” accounted for 70% of the population interaction diversity, on average.

Our results highlight how downscaling from species to individual-based networks helps understanding the structuring of interactions within ecological communities and provide an empirical basis for the extension of niche theory to complex mutualistic networks.

**Significance Statement:** Ecological interactions in nature occur between individual partners rather than species, and their outcomes determine fitness variation. By examining among-individual variation in interaction niches, we can bridge evolutionary and ecological perspectives to understand interaction biodiversity. This study investigates individual plant variation in frugivore assemblages worldwide, exploring how plant individuals “build” their interaction profiles with animal frugivores. The structure of networks composed of individuals was surprisingly similar to networks composed of species. Within populations, only a few plants played a key role in attracting a high diversity of frugivores, making them central to the overall network structure. Individuals actually interacted with a substantial diversity of partners, with individual niche “breadth” accounting for up to 70% of total interaction diversity, on average.

## Introduction

Species are a fundamental unit of study in most ecological research, resulting in numerous theoretical and methodological approaches to assess how their interactions support ecosystem functions. Network ecology based on graph theory has emerged as a useful framework to study these multi-species interactions simultaneously and assess the complexity of natural ecosystems (Solé & Valverde 2004, Fortuna & Bascompte 2008, Fontaine et al. 2011). Starting with food webs (Cohen 1978), network theory expanded its versatility to other ecological interaction modes such as mutualisms (Jordano 1987, Memmott 1999). Since then, abundant literature has revealed emergent and global properties of ecological networks, highlighting surprisingly similar architecture in the way they are assembled (McCann et al. 1998, Mora et al. 2018). Among ecological networks, mutualistic networks represent mutually beneficial interactions, and their structure and topology have been extensively explored (Bascompte & Jordano 2007). Plant–animal mutualistic networks are highly heterogeneous (i.e., most species have few interactions while a minority of species are much more connected) and nested (i.e., specialists interact with subsets of the species with which generalists interact), leading to asymmetric dependences among species (Jordano et al. 2003, Bascompte & Jordano 2007). Yet, it is not clear to what extent these properties emerge from networks at lower levels of organization, such as those composed of individual interactions.

Although interaction patterns are usually summarized at the species level, ecological interactions actually occur as encounters between individuals rather than species (Clark et al. 2011). For instance, while we say blackbirds (*Turdus merula*) consume fruits and disperse raspberry (*Rubus idaeus*) seeds, it’s actually individual plants and birds interacting within a local population. By missing this individual-level resolution we miss two important opportunities: 1) the ability to effectively link individual trait variation with interaction outcomes (fitness effects) and thus connect ecological and evolutionary perspectives; and 2) to bridge the gap between niche theory and complex interaction networks, i.e, to assess how individual-based interactions scale up into complex interaction networks.

Classic studies of animal-mediated seed dispersal interactions have been plant-focused (e.g., Snow & Snow 1988), and provide a useful framework to zoom-in into the interactions established between a particular plant species and a set of animal frugivores. By considering individual-based networks, in which one set of nodes is composed of plant individuals, and the other set is composed of animal species, we can examine individual variation in “interactions build-up”, as well as its subsequent implications, in e.g., fitness (Gómez & Perfectti 2012, Rodríguez-Rodríguez et al. 2017, Arroyo-Correa et al. 2021). This is helpful not just for building a proper bridge between interaction ecology and demographic consequences (e.g., Quintero et al. 2023), but also for bridging network ecology with evolutionary consequences (Guimarães et al. 2011, Segar et al. 2020).

Network structure may not be consistent across hierarchical scales of organization (Tur et al. 2014, Wang et al. 2021). The similarity in the set of partners available to individuals of the same species will be higher than that to different species. That is, the physical and phenological traits of conspecific individuals tend to be more similar than those among species (Siefert et al. 2015), discouraging major forbidden interactions (but see Albert et al. 2010, González-Varo & Traveset 2016) and likely increasing overall network connectance. Thus, we might expect individual-based networks to exhibit architectural and structural properties different to those found in species-based networks; yet, this remains an underexplored question.

Downscaling the study of interactions to individuals allows us to observe how the variation among individuals in their partner use is distributed in the population (Fig. 1A-B). Since its origins, the niche concept has provided an ideal framework for studying individual variation in resource use (Grinnell 1917, Van Valen 1965, Bolnick et al. 2003). Even so, most previous work has focused on antagonistic interactions such as predator–prey trophic niches (Bolnick et al. 2003, Araujo et al. 2011, Costa-Pereira et al. 2018, Costa-Pereira et al. 2019). It was only very recently that niche theory was applied for understanding individual variation in mutualistic interactions (Tur et al. 2014, Albrecht et al. 2018, Phillips et al. 2020, Koffel et al. 2021, Arroyo-Correa et al. 2023, Gómez et al. 2023). However, the study on how a plant individual’s interaction niche is distributed within a population remains largely unexplored, even less has been compared across different ecological systems. For this study, we rely on the concept of ‘interaction niche’ as the space defined by the set of species with which a population, or a plant individual, can interact (Fig. 1C) (Ponisio et al. 2019).

**Figure 1.**
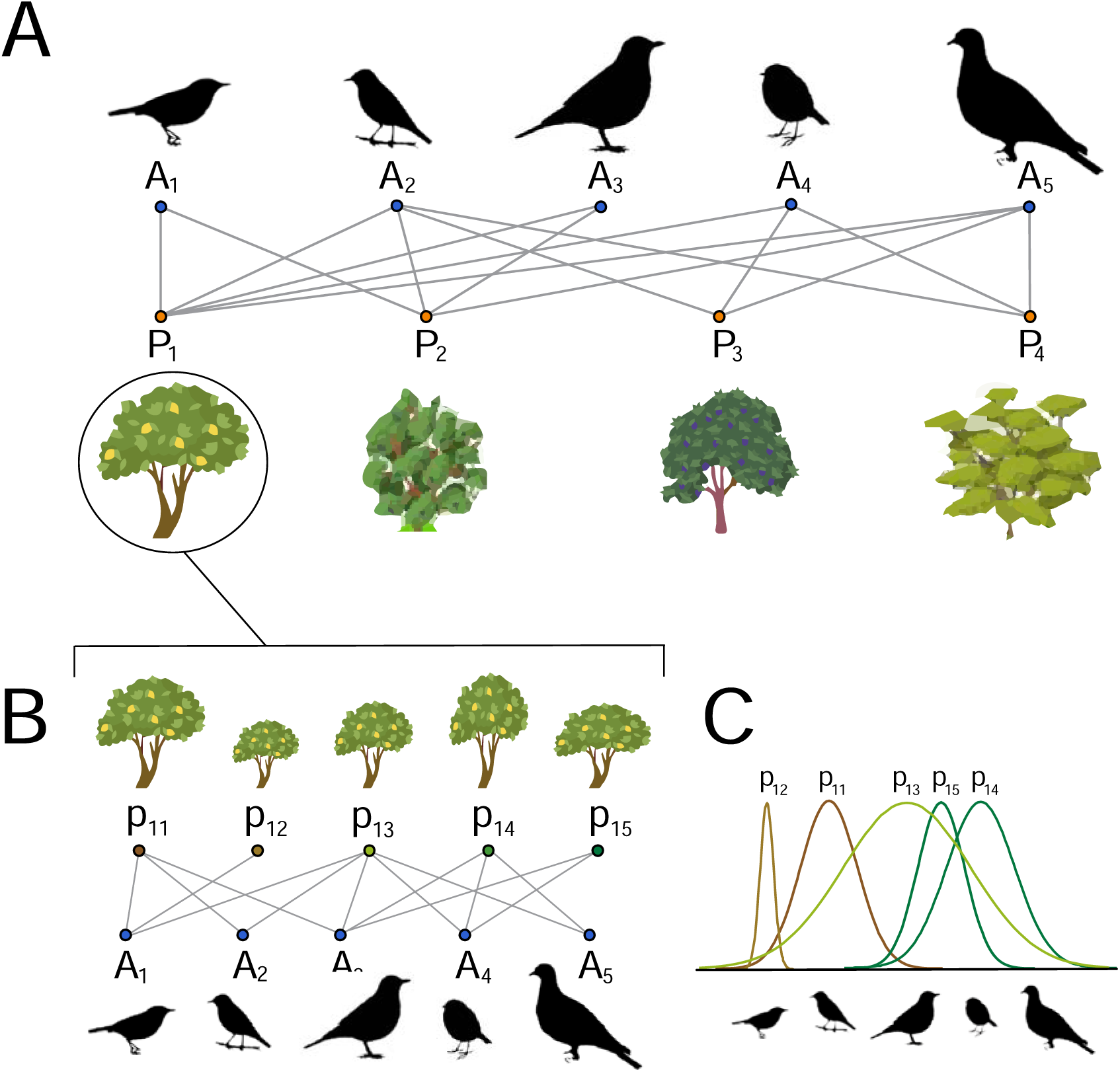
A) Schematic example of a species-based interaction network between four ornithochorous plant species (*P*_1_ - *P*_4_) and their frugivore assemblage with five animal species (*A*_1_ - *A*_5_) (top). B) A zoom in on the individual-based network of plant species P_1_ depicting the interactions of its plant individuals (*p*_11_ - *p*_15_) with five animal species, exemplifying the study focus of this paper. C) Different plant individuals (*p*_11_ - *p*_15_) interact with frugivore assemblages of variable diversity, illustrating their individual interaction niches. Plant niches are exemplified by the five colored niche utilization curves within the inset which indicate the relative interaction frequency with each of the

Interaction probabilities between plant individuals and animal species (i.e., probability of interspecific encounter, PIE; Hurlbert 1971, Chase & Knight 2013) are influenced by a myriad of factors such as population abundances, accessibility of resources, individual preferences or physiological needs (e.g., optimal foraging theory) as well as required matching in traits and phenology (Guimarães 2020). Intraspecific trait variability, neighborhood attributes and spatio-temporal context drive animal preferences for certain plant individuals, which will govern the establishment of interactions between plant individuals and their mutualistic partner species (Sallabanks 1993, Snell et al. 2019, Isla et al. 2023). In mutualistic systems such as pollination or seed dispersal, variation in the patterns of interaction or exploitation of niches (available partners) can play a determining role, as mutualists directly affect the reproductive outcome of individuals, influencing fitness variation and trait selection, which act as raw material for coevolution (Thompson 1999), as well as population dynamics and community assembly (Simmons et al. 2020, Arroyo-Correa et al. 2023).

Quantifying individual variation in interaction niche, and particularly niche partitioning, can shed light on the coexistence and stability of mutualistic communities. For instance, individuals in a population can behave as specialists or generalists when exploiting their interaction niche, and this may influence how these individuals are affected by interspecific competition and how partner diversity is promoted, determining, e.g., degree distributions in interaction networks (Bascompte & Jordano 2014). The extent to which individuals behave as specialists or generalists in a population can be elucidated by partitioning niche variation into its between-(BIC) and within-individual (WIC) components. Thus, this approach can prove useful to predict niche-shifts or interaction niche expansion (Roughgarden, 1972, Bolnick et al. 2007). The levels of individual specialization in individual-based networks can be estimated as the proportion of the total niche width in the population (TNW; total partner diversity) due to within-individual variation (WIC; average partner diversity of individuals). Thus, the distribution of frugivore–partner species richness and interaction allocation among plant individuals can be highly variable in local populations (e.g., Jordano & Schupp 2000, Guerra et al. 2017, Miguel et al. 2018, Jácome-Flores et al. 2020, Quintero et al. 2023). By studying plant individual specialization and how frugivores distribute interactions among plants, we aim to understand variation in mutualistic interactions within plant populations (Fig. 1). Examining how interaction niches are partitioned globally can expand the concept of niche variation to mutualistic interactions and pave the way for future hypotheses.

A variety of node-level metrics for complex networks can provide insight into an individual’s strategy within its population (Dormann 2011, Poisot 2012). Several studies have used node-level metrics to characterize individuals’ positioning in the network, informing us about their role and significance in their population (e.g., Gómez & Perfectti 2012, Guerra et al. 2017, Rodríguez-Rodríguez et al. 2017, Crestani et al. 2019, Vissoto et al. 2022, Isla et al. 2023, Arroyo-Correa et al. 2023). However, most of these studies have used a single or several metrics separately to understand the interaction profile of individuals and for single populations. By using a combination of node-level metrics, we aim to characterize interaction profiles of plant individuals with frugivore species and the distribution and frequencies of roles among and within populations in different geographic regions. Given contrasted differences in e.g., frugivore diversity and life histories across biogeographic realms, we could expect plant individuals from certain regions to exhibit similar interaction profiles, markedly different from those of individuals from other species and/or regions. Conversely, if life history or context-dependent effects were not determinant in structuring interactions between plant individuals and their frugivore partners, we could expect consistent individual interaction profiles across populations, irrespective of geographic location or biome type.

Obtaining an insightful picture of a mutualistic network structure from field data is a challenging task. When sampling is limited, the inferred network structure can be noisy, even biased, and thus subject to sampling fluctuations. This issue becomes particularly relevant when comparing networks from different studies (Brimacombe et al. 2023). Here, we build upon Young et al. (2021) Bayesian framework for reconstructing mutualistic networks to infer each pairwise interaction in individual-based networks, accounting for sampling completeness and the inherent stochasticity of field observation data. We then propagate the uncertainty of all pairwise interactions down through niche specialization and network interaction profiles.

The overarching goal of this study is to investigate the role played by individuals in the assembly, structure and functioning of complex ecological interaction networks. To do so, we combine network and niche theory to characterize the interaction profile of plant individuals in mutualistic seed dispersal systems across different bioregions aiming to illustrate the wide diversity of plant populations considered. We outline three main objectives: 1) examine whether networks composed of individuals exhibit different architectural and structural properties than those found in species-based networks, 2) understand how variation in frugivory interactions takes place at the plant population level by quantifying individual niche-partitioning and frugivore interaction allocation, and 3) characterize interaction profiles of plant individuals with frugivore species and assess the distribution and frequencies of roles among populations.

## Results

### Structure of individual versus species-based networks

We assembled a total of 46 individual-based plant-frugivore networks and compared them with 59 species-based networks using six network metrics (connectance, nestedness, modularity, assortativity, centralization and interaction evenness). Networks showed a remarkable overlap in all metrics at both resolution scales (Fig. 2A, Fig. S4, Table S3). Species-based and individual-based networks presented wide variation in their structure. Overall, when controlled by network relative size and interaction abundances, networks were less connected, nested and even, and more modular, assortative and central than their corresponding null models, although some of these differences were very subtle and consistent for networks at the two scales (Fig. 2B, Fig. S4). Comparing across scales, individual-based networks were slightly less connected and less modular than species-based networks relative to random expectations. However, these differences remained subtle and highly overlapping (e.g., 0.05 average difference in connectance and 0.04 in modularity; Table S3).

**Figure 2.**
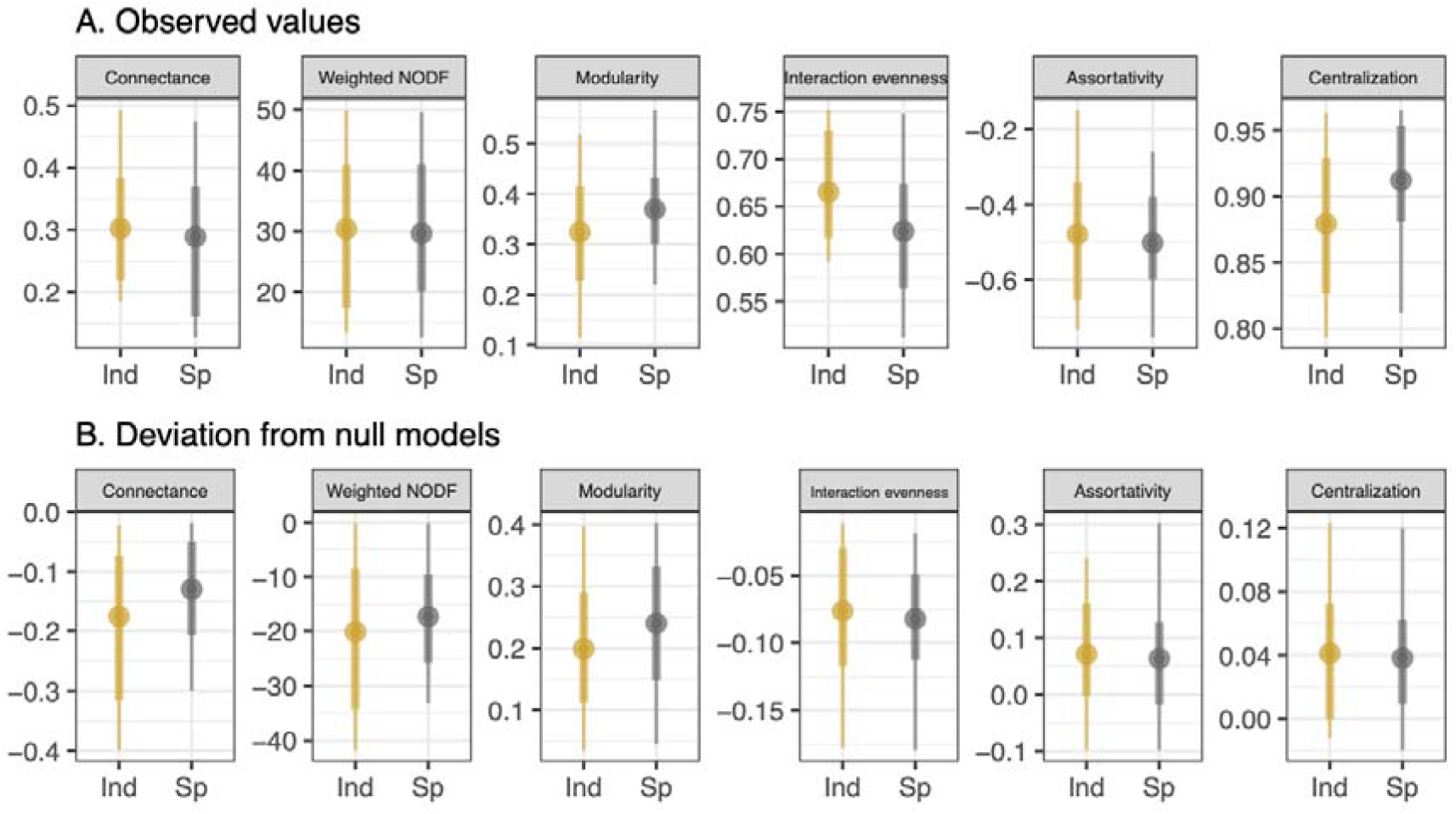
Network metrics comparison across network scales (Ind = individual-based, n = 46; and Sp = species-based networks, n = 59). Panel A shows observed network-level metrics and panel B deviation of observed metric values from their corresponding null model, which represents a network of the same size and relative abundances (generated using Patefield’s algorithm). Points represent the mean and thick and thin lines represent 0.6 and 0.9 confidence intervals respectively. For each network metric, deviation from observed represents the difference between the observed value and the null expectation.

### Plant individuals’ specialization in interaction niche

The 46 populations included a total of 1037 plant individuals: 373 individuals (from 9 species) from the Mediterranean (Iberian Peninsula), 389 (17 species) from Tropical regions in Asia and America, and 275 (3 species) from Southern Temperate regions (Southern Brazil and Argentina). Our aim in the compilation of individual-based networks was to showcase the diversity of environments and their associated variations; this analysis revealed major gaps in data coverage on the African continent and in the Australasian region (see Table S1), so a proper comparison among biogeographic areas awaits more complete data. Most plant populations studied presented low to medium levels of individual specialization (mean WIC/TNW = 0.68; 90% CI = 0.32 - 0.95). Individual-based networks from the Mediterranean region showed a higher proportion of generalized plant individuals (Fig. 3; mean WIC/TNW = 0.82), whereas southern temperate and tropical populations presented higher levels of individual specialization (mean WIC/TNW = 0.56 and 0.65 respectively, p-value < 0.01). Plant populations interacting with higher numbers of frugivore species had a wider interaction niche (TNW, i.e., Shannon diversity index, r = 0.53, p < 0.01), but not necessarily higher levels of individual specialization (WIC/TNW) (Fig. S5). The degree of individual specialization did not increase as TNW increased, because WIC increased in the same proportion as TNW (slope of log-log model = 1.000, SE = 0.005). That is, differences in the degree of individual specialization between bioregions were achieved via changes in both the BIC and the WIC components.

**Figure 3.**
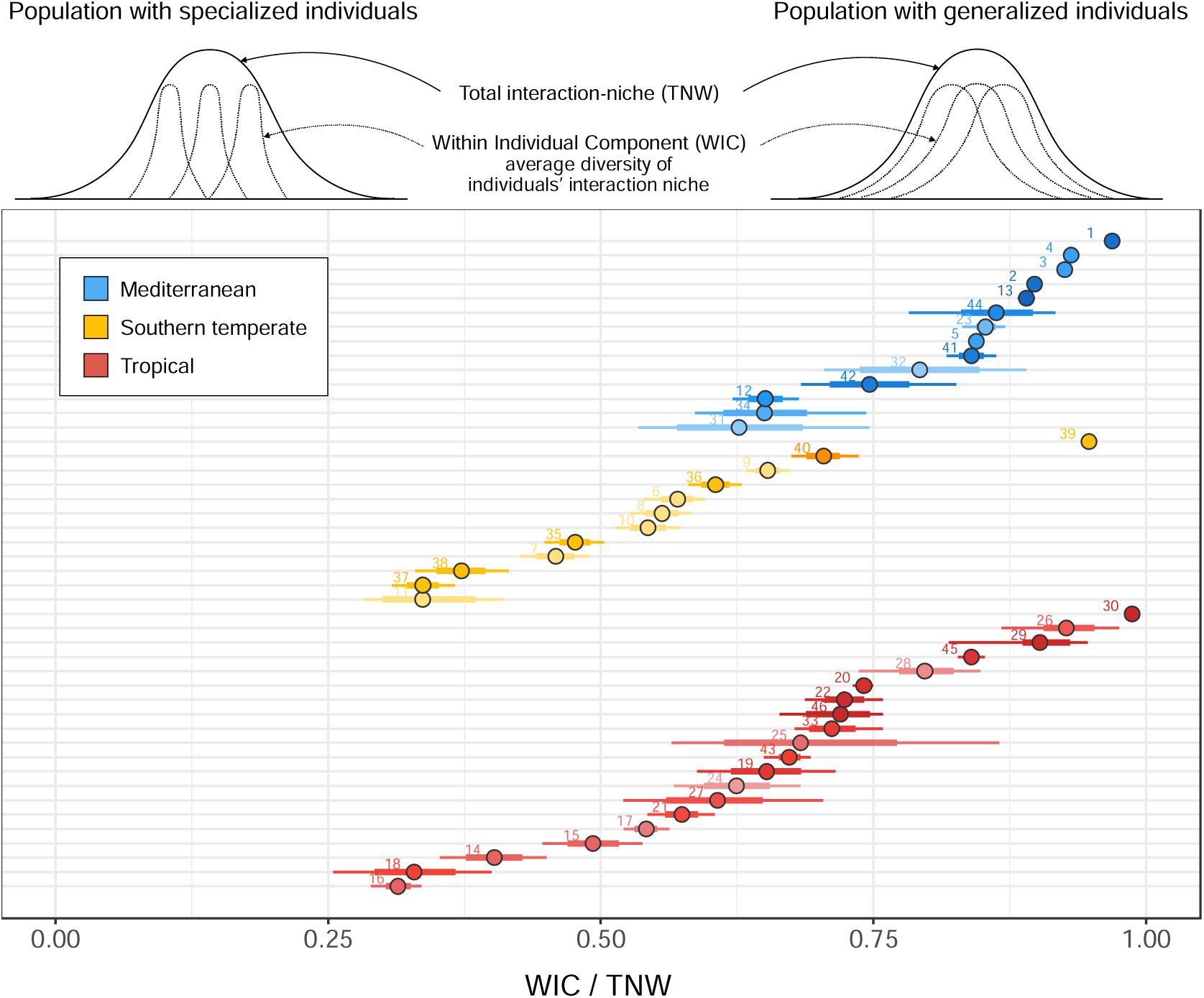
A schematic representation of two plant populations (top): the one on the left presents a population composed of highly specialized individuals (low WIC; low WIC/TNW) and the one on the right presents a population composed of highly generalized individuals (high WIC; high WIC/TNW). Values of WIC/TNW closer to 1 represent populations with generalized individuals where plants use most of the available interaction niche. On the other extreme, values closer to 0 indicate populations with specialized individuals that use a smaller subset of the available interaction niche (in this case plants do not tend to interact with the same frugivore species). Bottom panel: Values of individual specialization (WIC/TNW) for all plant populations studied (n = 46 networks). The total niche width (TNW) represents the interaction niche of the population, calculated as the Shannon diversity of interactions with frugivore species at the population level, i.e., aggregating across individuals. The within-individual component (WIC) is the average Shannon diversity of interactions with frugivores found within individual plants. Each point-interval in the graph represents an individual-based network (i.e., population), where the point indicates the mean WIC/TNW, and the thick and thin lines span the 60% and 90% credibility interval, respectively of each network posterior distribution. Colors represent the bioregions of the study site (see Table S1 for each network metadata).

### Frugivore interactions within plant populations

Across all plant populations and regions, we found that just a reduced subset of frugivore species (generally between one and three) usually accumulated most of the interactions, while the rest of frugivore species contributed a minor proportion. On average, more than half of these interactions were contributed by less than 20% of frugivores (SD = 8.2%), regardless of the total number of frugivore species in the plant population (Fig. S6). The frugivores that contributed most interactions also tended to interact with a higher number of plant individuals (Spearman’s rho = 0.81, Fig. 4). Remarkably, frugivore species with smaller contributions interacted with a variable proportion of plant individuals, such proportion being higher in Mediterranean networks and lower in Southern Temperate networks (Fig. 4). Frugivores’ body mass was not correlated with interaction contribution (rho = -0.08) nor with the proportion of plant individuals they interacted with (rho = - 0.13).

**Figure 4.**
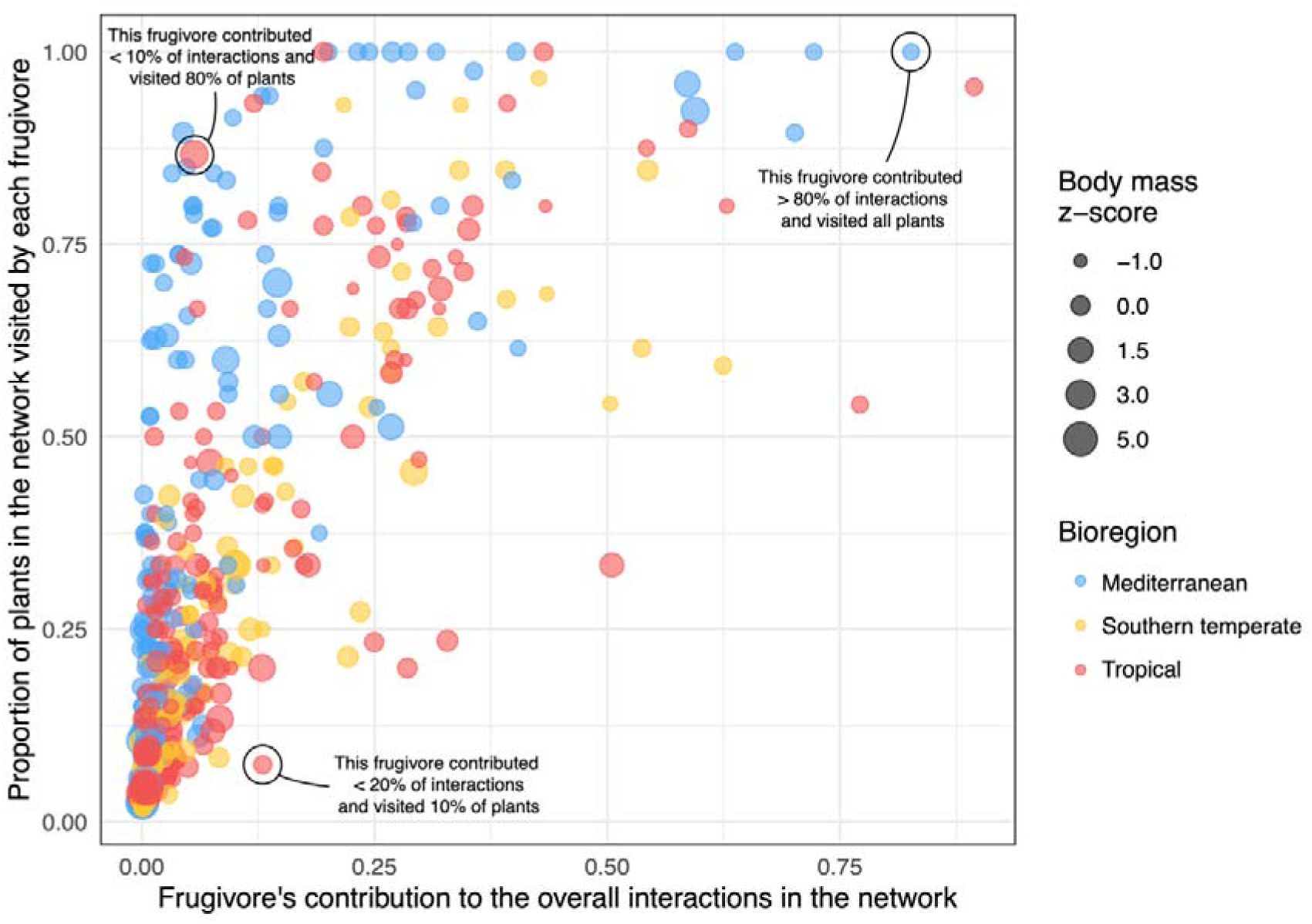
Relationship between the overall contribution to the total number of interactions by each frugivore species (i.e., link weights in the individual-based networks) and the proportion of plant individuals with which they interacted (i.e., animal degree). Each point represents a frugivore species in their respective plant population (individual-based network). Point color indicates the bioregion and point size is proportional to the frugivore’s body mass relative to other frugivores present in its population (z-score).

### Plant individuals’ interaction profiles

The multidimensional principal component analysis space occupied by all plant individuals and defined by node-level metrics (normalized degree, strength, specificity, overlap and weighted closeness) did not produce distinct clusters by bioregion or population. Instead, individuals from different populations spread across the multidimensional space, suggesting ample within-population heterogeneity in plant individuals’ interaction profiles (Fig. 5, Fig. S8). The first principal component (PC1), explaining more than half of the variation (51%), was mainly related to interaction degree and specificity, thus capturing individual variation in frugivore richness and composition of the plant individual’s assemblages. The second component (PC2) explained 24% of the variation and was correlated with niche overlap and interaction strength; these metrics are related to plant individuals’ interaction patterns in relation to their conspecifics and affected by interaction frequency (link weight). Plant individuals with more unique frugivore assemblages were positioned in the upper area of the PCA space, while many plants with highly-overlapping frugivore assemblages within their populations were positioned towards the bottom area. The third component (PC3; 10% variation explained; Table S4) was strongly related to weighted closeness, a measure of how strongly and well connected (i.e., central) individuals are within the network.

**Figure 5.**
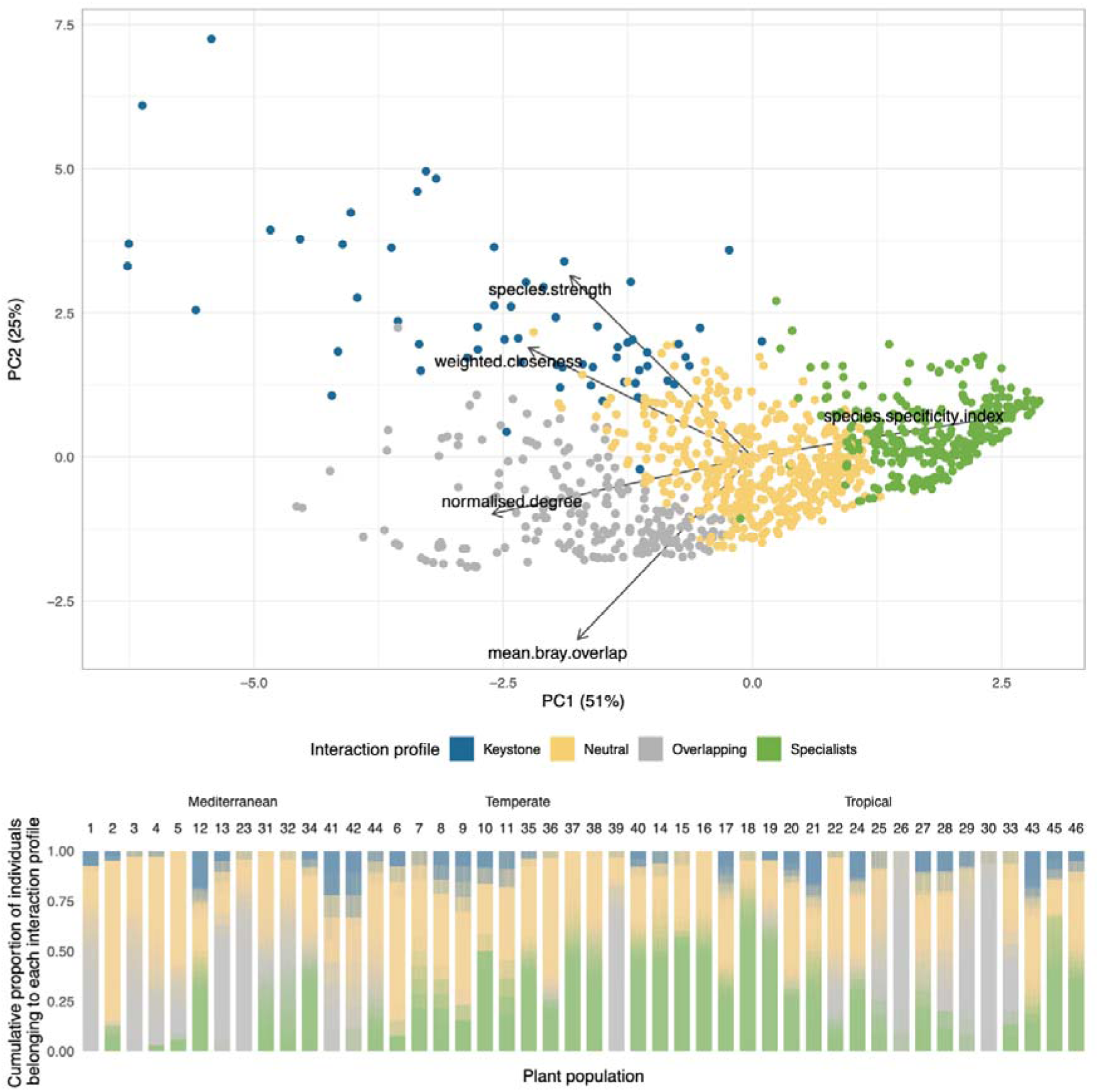
Principal Component Analysis for node-level metrics of plant individuals in all plant populations with the most common cluster attribution for each individual depicted in color. Each plant individual (points) is represented by the centroid of its 1000 node-metric values calculated with the full posterior distribution (Fig. S8). The panel below shows different plant populations as bars with a number code above (see Table S1). Each stacked bar is composed of 1000 thinner stacked bars that represent the cumulative proportion of individuals attributed to each cluster or interaction profile originated from the 1000 reanalysis using the posterior samples.

Few plants were highly central in the interaction network (high weighted closeness) and important for frugivore dependence (high species strength) (i.e., points in the upper-left area of the multivariate space). Most plant individuals showed uneven dependencies on frugivore species and/or medium-high frugivore overlap with other plants in the population. Yet plants with strong dependencies on one or few frugivore species tended to show lower overlap with other individuals in the frugivore assemblage, suggesting a trade-off between partner specialization and partner sharing (lower-right Fig. 5). Overall, plant individuals from the Mediterranean tended to have more similar frugivore assemblages (higher niche overlap), while plant individuals from Southern Temperate regions presented less overlapping and more specialized frugivore assemblages (Fig. S9).

A cluster analysis further revealed consistent interaction profiles within plant populations, regardless of bioregion or species. All networks presented a variable but substantial proportion (mean = 42%; 90% CI = 8 - 77%) of plant individuals with “neutral” interaction profiles, which did not excel in any node-level metric (Fig. 5, Fig. S9). The clusters “specialized” (mean = 28%; 90% CI = 0 - 67%) and “overlapping” (mean = 22%; 90% CI = 0 - 86%) included a variable but substantial proportion of plants, most likely reflecting either distinct frugivore assemblages (e.g., in the tropical biomes) or more overlapping in composition (mostly non-tropical), respectively. Notably, in most populations (36 out of 46), we detected a low proportion (mean = 8%, 90% CI = 0 - 24%) of “keystone” individuals showing high strength and weighted closeness. We termed these plant individuals as keystone since they accounted for a high proportion of interactions within their respective populations, playing a central role in plant-frugivore interaction structuring.

## Discussion

Our study highlights how downscaling from species to individuals uncovers consistent network structures across biological levels, mutualistic partner allocation among plants globally, and interaction profile similarities within populations, regardless of species or ecological context. This reveals new aspects of ecological interactions and network assembly at the individual level.

### Effects of downscaling resolution on network structure

The structure of plant–animal mutualistic networks revealed fundamental heterogeneity across networks and resolution scales. We did not find major deviations in the assembly patterns of interactions as we zoomed in from species to an individual-based scale. Previous research exploring the consequences of downscaling on network architecture found significant shifts in the structure of pollination networks (Tur el al. 2014, Wang et al. 2021): individual-based networks were less connected and individuals were more specialized than species. These studies examined one community using a single methodology. Our approach, however, involved diverse communities and populations from different regions with varying methodologies, capturing broader interaction patterns.

Surprisingly, we found no significant differences in network connectance across scales, contrary to our initial expectations. The slightly larger connectance of individual networks after Bayesian modeling (average connectance = 0.4 ± 0.2 vs 0.3 ± 0.1 in raw networks) suggests that undersampled interaction matrices could be missing interactions. On the other hand, the slightly lower modularity in individual-based networks could stem from a decrease in the number of forbidden links as frugivores can potentially interact with virtually all the plant individuals within populations (see González-Varo & Traveset 2016). In contrast,species-based networks involve much more heterogeneous sets of plant species. The addition of new species or individuals with new traits provides new link possibilities in a network, yet in the case of species, potential interactions must undergo stronger trait and phenological matching filters than in individual-based networks (Sazatornil et al. 2016). Simply put, a given frugivore species may interact with a broader range of partners within a plant population than when interacting with the full range of taxonomically-diverse available plant species in a community. The former set imposes less constraints to interactions by including much more homogenous conspecific partners. In this way, downscaling from species to individuals fundamentally alters the probabilistic distribution of interactions among partners (Poisot et al. 2016).

Aside from minor differences in certain network metrics, the overall topology and structure of frugivory networks at different resolution scales were not sufficient to make clear distinctions. This convergence in network structure may be driven by factors like the probability distribution of interspecific encounters (PIEs), influencing network configurations consistently across scales. We argue that numerical effects are likely at the base of these emergent properties, governing interactions distribution across nodes and asymmetric interactions (Jordano 1987, Vázquez et al. 2007, Schleuning et al. 2011, Guimarães 2020). These numerical effects can be caused by varied organism abundances in the case of species, or traits and genotypes in the case of individuals, that modulate the attractiveness of plants to frugivores, such as crop size, plant height, or phenology (Snell et al.

### Individual specialization in the interaction niche

Individuals’ interaction niches were narrower than those of their populations, supporting that individual specialization is substantial and common in nature (Bolnick et al. 2003), even in mutualisms. Plant individuals’ specialization levels were similar to levels reported in other animal taxa (Araujo et al. 2011). Interestingly, the degree of individual specialization varied across biogeographic regions, yet most plant species showed WIC/TNW ratios >50%, which indicates moderate generalization among plant individuals. Broader and more overlapping frugivore assemblages in Mediterranean regions versus higher specialization and variability in southern temperate and tropical networks could not be attributed to differences in taxonomic diversity as all regions presented similar total niche width (Fig. S5). Instead, plant individuals in southern temperate regions presented smaller relative niches. Tropical populations consisted of frugivore assemblages of variable diversity and highly variable individual specialization, yet limited area coverage of the available individual-based networks hindered comparisons. No significant niche breadth differences were found across bioregions, aligning with studies of terrestrial food webs, challenging the latitude-niche breadth hypothesis that predicts narrower niches in tropical regions (Cirtwill et al. 2015, although see Araujo & Costa-Pereira 2013). The large variation in individual specialization within bioregions may be pointing to the role of fine-scale factors such as the local habitat, neighborhood effects, or the influences of individual phenological variation. Further research is needed to evaluate the ecological correlates of plant individual interaction niche utilization and its consequences.

Different levels of individual specialization can have implications for population stability (Arroyo-Correa et al. 2024) and niche expansion (Van Valen 1965). According to the niche variation hypothesis, populations experiencing niche expansion achieve it through increasing their inter-individual variation (Bolnick et al. 2007). By diversifying its resources, plant individuals would be able to exploit novel and underutilized frugivores escaping competition from conspecifics. Niche shifts and expansion have become exceptionally important for adaptation to changing climate conditions (Hallfors et al 2023) as well as changes in frugivore assemblages and fluctuating abundances (Campos-Celada et al. 2022). Therefore, the variation we found among populations in frugivore assemblage specialization will likely have an impact on the adaptation of plant–frugivore mutualistic interaction niche in current and future scenarios of global change.

In all plant populations just a few frugivore species, even within diversified assemblages, consistently perform most of the mutualistic interactions (Fig. S6; Rother et al. 2016, Guerra et al. 2017, Isla et al. 2023, Rehling et al. 2023, Thiel et al. 2023). Although frugivore body mass did not prove to be a good indicator of their contribution to interactions (although see Valenzuela-Ospina & Kattan 2021), it may play a role in seed dispersal effectiveness due to its positive correlation with the number of fruits consumed per visit or the frequency of long-distance seed dispersal events (Snow & Snow 1988, Jordano et al. 2007, Godínez-Álvarez et al. 2020). These highly uneven interaction patterns will result in asymmetric dependencies between plant individuals and frugivore species, where the main frugivore shows low specificity for specific plants, while most plant individuals rely mostly on the main frugivores’ service (Quintero et al. 2023). Asymmetric dependency between partners also emerges at species–species interaction level (Jordano 1987, Vázquez & Aizen 2014, Bascompte et al. 2006); further downscaling into individual-individual interactions would help elucidate if asymmetry remains consistent across scales. Finally, our analysis reveals a consistent trend for frugivory and seed dispersal service in a given plant population (estimated from the proportion of plant individuals with which a frugivore species interacts) to increase with the overall contribution to the total number of interactions by each frugivore species (i.e., link weights in the individual-based networks, Fig. 4). Thus, central frugivores interact with a wide range of plant individuals, most likely an emergent result of the interaction asymmetry discussed above.

### Consistency of individual plants’ interaction profiles across regions and populations

Plant individuals’ interaction profiles were not explained by bioregion or species, pointing to fundamental architectural patterns in the assemblage of mutualistic interactions that are not strongly constrained by phylogeny or geographic location but rather by the interplay between traits and numerical effects (Jordano 1987, Carnicer et al. 2009, Albrecht et al. 2018, Guimarães 2020). Remarkably, we found a consistent distribution of plant interaction profiles within populations, with most individuals acting in an average manner, a variable fraction standing out for their specialization or redundancy and only very few individuals having a central role, high diversity of interactions, and strong frugivores’ dependence on them (“keystone” plant individuals). Similar results are reported in food webs, where a core group of species shares ecological roles, while peripheral species have unique interaction profiles (Mora et al. 2018). It is likely that within frugivory networks these key individuals present unique phenotypic traits, such as abundant fruit crops or advantageous locations that make them reliable to many frugivores (Snell et al. 2019, Isla et al. 2023).

Although some of the plant species considered in this study were generalists within their community, individuals in their populations showed variable interaction niche breadths (Guerra et al. 2017), with populations consisting of both generalist and specialist individuals (Arroyo-Correa et al. 2023, Bolnick et al. 2007). This mix creates species that appear broadly interactive but actually include individuals with varied interaction patterns, from wide generalization to specific partner preferences. This highlights the complexity of species’ ecological roles within communities. (Guimarães 2020).

### Concluding remarks

We found consistent patterns of interaction assembly across biological scales using a set of biologically informative network metrics. On top of the absence of a clear hierarchy differentiation in network structure between individuals and species, we found that almost every individual-based network included a similar representation of individual interaction profiles, evidencing a common backbone in the way interactions are organized (Mora et al. 2018). Conducting future analyses on interaction types or motifs of individual-based networks may provide us with new insights, as these approaches have proven effective in distinguishing networks between and within ecological systems (Mora et al. 2018, Michalska-Smith et al. 2022, Pichon et al. 2023).

Intraspecific variation appears as a central ingredient in the configuration of complex networks of mutualistic interactions, driven by the widespread interaction profiles of frugivore species with plant individuals. High levels of intraspecific variation have been shown to confer greater stability to mutualistic systems (Arroyo-Correa et al. 2023). By zooming in on ecological interactions this study provides valuable insights into how mutualistic interactions are similarly structured at the individual level and reveal underlying, consistent, patterns of role assignment within populations and across bioregions.

## Methods

### Dataset acquisition

We compiled frugivory ecological networks , both at the species and the plant individual level. Species-based networks were gathered from 40 published studies at the community scale (see Table S1). For individual-based networks, which are scarcer, we compiled phyto-centric studies (plant-based), with quantitative information on frugivore visitation on plant individuals within populations. We combined published studies with unpublished datasets, gathering data for 21 different study systems, including datasets from our own field studies with different Mediterranean species (n = 9). Some of the studies selected presented more than one network from different communities (in species-based studies) or populations (in individual-based studies) (Table S1).

These datasets document interactions between plant species and animal frugivore species (in species-based studies) or interactions between plant individuals of a single species that coexist in a local population and animal frugivore species (in individual-based studies). None of the datasets collected are nested, that is, individual-based networks are not sampled within the same study as a species-based network, which prevents a direct structure dependence between the two scales of the datasets. Data was entered as adjacency matrices, where rows represent plant species (or individuals) and columns represent animal species, with matrix elements *a*_ij_ indicating interaction frequency (visitation frequency to plants). In order to ensure networks were sufficiently sampled to robustly characterize their structure and interaction profiles, we only kept those that were reasonably complete. We checked for sampling coverage of individual-based networks using iNext R package (Hsieh et al. 2016) (Table S2). To do so, we converted the adjacency matrix data to an incidence frequency-data and considered plant individuals as sampling units and the number of frugivore species detected at each plant (species richness). We discarded networks in which the number of interacting nodes (plants and frugivore species) was less than 15 or plants were less than six (n = 13 networks). Our final dataset consisted of 105 networks with an average size of 384 potential links or cells (range = 65 - 2904) and 90 unique interactions (range = 22 - 419). Forty-six were individual-based networks and 59 were species-based networks (Table S1). When possible we referred the interaction value to the coarsest level, that is, frugivore visitation events, otherwise number of fruits consumed.

### Network-level metrics

For both the individual and species-based networks, we calculated several network-level metrics, using R packages bipartite (Dormann et al. 2008) and igraph (Csárdi & Nepusz 2006). All network metrics were calculated using standardized matrices to minimize the impact of differences in sampling effort and study-site characteristics. Specifically, for all those individual-based networks where sampling effort across plant individuals was heterogeneous within the population (21 out of 46 networks), we divided plant visits by the amount of time observed and/or the area sampled in each plant, so that interaction counts were comparable. Subsequently, all networks (individual and species-based) were scaled by dividing the weight of each pairwise interaction by the total number of interactions in the matrix (grand total standardization; Quintero et al. 2022). In this way, the interaction values (matrix cells) represent the relative frequency of a plant individual (in individual-based networks) or a plant species (in species-based networks) interacting with a given frugivore species, and the sum of all relative frequencies equals one.

We selected a representative set of metrics that had suitable biological interpretation and were not highly correlated (Variance Inflation Factor < 3) and/or not strongly affected by the number of species/individuals sampled or overall network size (Fig. S1, S2, S3).

Selected network-level metrics were:

1. Connectance *(topology).*This metric gives the proportion of realized over potential links in the network. Calculated as the sum of realized links (unweighted) divided by the number of cells in the matrix. Values range from 0 (no links) to 1 (fully connected networks where all nodes interact among them) (Dunne et al. 2002).
2. Weighted nestedness wNODF *(structure)*. It informs on the way interactions are organized. A highly nested structure is one in which nodes with fewer connections tend to interact with a subset of highly connected nodes that in turn interact with the highly connected ones (Bascompte et al. 2003). Values of 0 indicate non-nestedness, those of 100 perfect nesting (Almeida-Neto & Ulrich 2011).
3. Assortativity *(topology)*. This metric indicates the level of homophily among nodes in the graph. It ranges from -1 to 1, when high it means that nodes tend to connect with nodes of similar degree; when low, nodes of low-degree connect with nodes of high-degree (disassortative) (Newman 2002, Barabasi 2016).
4. Modularity *(structure - clustering*.)It reflects the tendency of a network to be organized in distinct clusters or modules (Bascompte & Jordano 2014). This metric ranges from 0 (no clusters) to 1, where nodes within a module interact more among them than with nodes from other modules (highly compartmentalized network; Newman 2006).
5. Eigenvector centralization *(centrality)*. This metric quantifies how centralized or decentralized the distribution of eigenvector centrality scores is across all nodes in a network (Freeman et al. 1979). The eigenvector centrality of a given node in a network is a measure of the influence of that node, taking into account both the node’s direct connections and the connections of its neighbors. Nodes with high eigenvector centrality are connected to other nodes that are also central, giving them a higher score (de Oliveira Lima et al. 2020). The network-level eigenvector centralization provides a measure of the extent to which a few nodes dominate the network in terms of influence. In a network with low centralization, the centrality scores are relatively evenly distributed among the nodes, suggesting a more decentralized structure where many nodes contribute to the overall connectivity of the network, and therefore to the interaction services. On the other hand, a network with high centralization indicates that only a small number of nodes have a higher centrality, suggesting a more centralized structure where a few nodes play a crucial role in the network’s overall connectivity. We normalized this measure to ensure that the centralization value is relative to the maximum centralization for a network of a given size.
6. Interaction evenness (*interaction diversity*). Also known as Pielou’s evenness, it quantifies how balanced the distribution of interactions is across nodes (Blüthgen et al. 2008). This metric considers all links as species, and their weight as a measure of their abundance. It quantifies link diversity using Shannon index and divides it by the theoretical maximum richness of all possible links (i.e., logarithm of Shannon). It ranges from 0 to 1, where low values indicate high unevenness and high values approach perfect evenness in interaction distribution.

### Comparison across network scales

We used the aforementioned metrics to compare the structure and assemblage of species-based versus individual-based networks. Additionally, to account for network size and inherent characteristics of each study, we assessed metric deviations against 1000 randomizations using null models. We use a Patefield algorithm (Patefield 1981, Dormann et al. 2009) that maintains network marginal totals. For each network and metric, we calculated the difference between the observed values and the randomizations. This difference indicates both the magnitude and direction in which observed values deviate from what could be expected at random.

### Bayesian modeling of individual-based network structure

Reconstructing mutualistic network structure from field data is a challenging task. Interaction data are hard to collect and typically require large sampling efforts, particularly to characterize infrequent interactions. Inferred network structure is highly sensitive to sampling design, effort, and completeness (Jordano 2016). Comparing networks from different studies without accounting for these sampling effects may lead to mistaken inferences (Brimacombe et al. 2023). Here, we build upon Young et al. (2021) Bayesian framework for reconstructing mutualistic networks to infer the posterior probability of each pairwise interaction in individual-based networks, accounting for sampling completeness and the inherent stochasticity of field observation data.

Following Young et al. (2021), pairwise interaction counts between plants and animals can be modeled as following a Poisson distribution whose mean (*μ*_ij_) is determined by the sampling effort spent on each plant (*C*_i_), the relative interaction abundance of each plant and animal in the population (*σ*_i_ and *τ*_j_), the (inferred) existence of an interaction link among both partners (*B*_ij_), and the “preference” parameter *r* which represents the difference in the average number of visits or interactions when there is a connection between mutualistic partners (i.e., when a frugivory preferentially interacts with a given plant, *B*_ij_ = 1) compared to when there is no connection (*B*_ij_ = 0).

Thus,

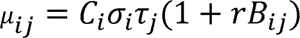

The preference parameter *r* accommodates the expectation that the number of interactions will be higher when a frugivore prefers a given plant. When there is no preference the number of interactions can still be higher than zero, but lower in principle. Here we adapted the model proposed by Young et al. (2021) to allow for varying preferences (i.e., interaction counts) among different frugivore species, so that some frugivores may bring many more visits than other mutualists. Rather than having a single preference parameter *r* for all species, each frugivore has their own preference *r*_j_ which is drawn from an exponential distribution with rate parameter beta = 0.01. In order to facilitate modeling and incorporate these modifications, we developed an R package called ‘BayesianWebs’ (Rodríguez-Sánchez 2024a) which relies on Stan (Stan Development Team 2024) and cmdstanr (Gabry et al. 2024) for parameter estimation, and contains functions to facilitate modeling of bipartite mutualistic networks including data preparation, model fitting, checking and visualization. Using this package, we fitted a Bayesian model to each individual-based network. As model output, we obtained 1000 posterior samples of the expected count for each pairwise interaction. Posterior predictive checks showed the expected increase of uncertainty in networks with limited sampling, and overall confirmed the good match between the observed data and predicted counts.

To ensure comparability across networks, posterior interaction counts were rounded to integer values and then standardized following the same procedure explained above. Networks with heterogeneous sampling effort across plant individuals were adjusted by dividing the posterior counts by the individual relative observation time or sampled area. Subsequently, all networks were standardization, Quintero et al. 2022), so that matrix cell values reflected the relative frequency of each interaction within the population.

The resulting distributions of pairwise interaction counts generated by the Bayesian model for each network (1000 matrices for each network or population) were then propagated all the way down to niche specialization and individual node-level metrics (see below).

### Niche specialization

We estimated populations’ niche specialization using the Shannon approximation of the WIC/TNW index for discrete data (Roughgarden 1979, Bolnick et al. 2002) implemented in the ‘network.tools’ R package (Rodríguez-Sánchez 2024b). In this case, we define as a niche-resource the available coterie of visiting frugivore species in a given population. This index computes the relative degree of individual specialization as the proportion of total niche width (TNW) explained by within-individual variation (WIC). Total niche width (TNW) is calculated as the total diversity of frugivore species visiting the plant population, using Shannon index,

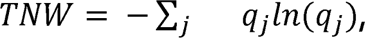

where *j* represents an animal species, and *q*_j_ is the proportion of interactions contributed by frugivore species *j* to the total number of interactions in the plant population.

The within-individual variation (WIC) is calculated as the average Shannon diversity of frugivores for each plant individual, weighted by the relative proportion of all frugivore interactions in the population that are used by each individual,

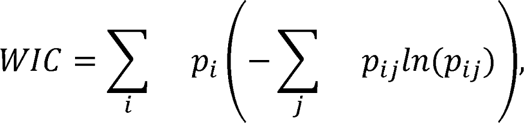

where *i* presents a plant individual, *p*_i_ is the proportion of interactions contributed by plant *i* to the total number of interactions in the population, and *p*_ij_ the proportion of interactions that animal species *j* contributed to plant individual *i*.

Finally, WIC is divided by TNW. Values closer to 1 indicate a population composed of generalist individuals that are using most of the niche available to the population. On the contrary, values closer to 0 indicate a population of specialist individuals using small subsets of the population niche, with large differences in resource-use among them. WIC and TNW estimates were calculated for each of the posterior counts (n = 1000) generated from Bayesian models for each network, rendering a credibility interval of individual specialization for each plant population. To test differences in individual specialization (i.e., WIC/TNW) between different bioregions (n = 3) we fitted a mixed-effects linear model with a Normal distribution where the network and study were present as random factors (Bates et al. 2015).

### Node-level metrics

To characterize plant individuals’ interaction profiles in their populations, we computed a set of node-level indices for each plant individual using R package bipartite (Dormann et al. 2008).

Additionally, we calculated average niche overlap using Bray-Curtis dissimilarity index (vegan R package; Oksanen et al. 2022). We computed these metrics using the full posterior distribution of each individual-based network. That is, for each of the 1000 posterior matrices generated for each network we calculated node-level metrics, therefore we obtained 1000 values for each plant individual and metric. We selected a representative set of metrics that had suitable biological interpretation for assessing the individuals’ interaction profiles and were not highly correlated nor affected by the number of individuals sampled (Fig. S7).

Selected node-level metrics were:

1. Normalized degree *(interaction diversity)*I.t represents the richness of partners for a given node and is scaled relative to the rest of nodes in the network. This metric ranges from 0 to 1, where a plant individual would score 1 if it interacts with all the frugivore species available (Dormann 2011).
2. Species specificity index *(interaction diversity)*. This metric informs about the variation in the distribution of interactions with frugivore species partners. It is estimated as a coefficient of variation of interactions for each plant individual, normalized to range between 0 and 1 (Julliard et al. 2006, Poisot et al. 2012). High values indicate higher variation in dependence on frugivore species. Plants with high dependence on few or a single frugivore species yield values close to 1, and plants that distribute their interactions equally with many frugivore species show low values.
3. Species strength *(interaction intensity).* It quantifies the dependence of the community on a given node (Dormann 2011), in this case, the frugivores represent the community and the plant individuals the nodes. It is calculated as the sum of the dependencies of each frugivore species (i.e., the fraction of all visits to a given plant individual coming from a particular frugivore species) (Bascompte et al. 2006).
4. Weighted closeness *(node position).* This metric provides an index of the magnitude to which a given node has short connection paths to all other nodes in the network (Opsahl et al. 2010). It is influenced by the intensity and number of links and indicates to what extent a node is in the “center” of the connections of the graph. This metric is calculated on the unipartite projection of the individual-based network for the plant individuals, with links between plant individuals representing the number of frugivore interactions shared. The weighted closeness of a plant individual is estimated as the inverse of the sum of all path lengths (link weights) between this plant individual and all other plant individuals in the unipartite network. Individuals with higher values of weighted closeness are strongly connected with more plant individuals in the population through shared frugivore species.
5. Mean interaction overlap using Bray-Curtis index *(node similarity).* This measure of interaction overlap informs on the average similarity in frugivore use between pairs of plant individuals. This metric indicates how different the frugivore assemblage of a given plant individual is compared to the rest of the population (e.g., Gómez et al. 2010). Higher values (i.e., higher overlap) indicate a higher similarity in interaction assemblage for a given plant individual with respect to other individuals in the population.

### Comparison across individual-based networks

In order to determine variation distribution in interaction structuring and node topology we performed a principal component analysis (PCA). Previous studies have used PCA for comparing network metrics (e.g., Sazima et al. 2010, Mora et al. 2018, Medeiros et al. 2018, Burin et al. 2021, Acevedo-Quintero et al 2023). In order to compare plant individuals’ interaction profiles, we performed a PCA using the full set of node-level metric distributions estimated for plant individuals within their population (i.e., individual node metrics derived from the full Bayesian posterior of each individual). This resulted in an ordination of elements (1000 points per each plant individual, single values sampled from the full posterior distributions of each individual-based network) in relation to the multivariate space defined by node-level metrics. Thus, such PCA provides an exploratory analysis of how plant individuals span the multivariate space of network metrics, where the location of each individual characterizes its interaction profile.

With the aim of identifying distinct plant individual profiles we performed a discriminant analysis of principal components (DAPC) using the adegenet R package (Jombart et al. 2023). We performed 1000 DAPCs using node-level metrics derived from the 1000 posterior samples of the individual-based networks. We selected four groups or clusters as the minimal number that would facilitate interpretation of each plant individual performance given its position in the network. In each of the 1000 analysis runs, plant individuals were assigned to one of the four groups given the highest probability provided by the DAPC. This analysis was based on the variation captured by the first 3 PCs which together explained 86% of the variance, with eigenvalues >0.7. Cluster groups were named post-hoc according to plants’ relative position in the PCA into ‘Keystone’, ‘Overlapping’, ‘Specialist’, and ‘Neutral’ interaction profile types. We subsequently quantified the percentage of individuals assigned to these four groups for each plant population in each of the 1000 repetitions.

### Data availability and code

All datasets and code used to generate this study are available for download in Zenodo (https://doi.org/10.5281/zenodo.10630818; Quintero et al. 2024) and GitHub repository (https://github.com/PJordano-Lab/MS_individual-based_networks).

## Acknowledgments

We are grateful to Miguel Jácome, Gemma Calvo, Pablo Homet, Pablo Villaba, Juan Miguel Arroyo, Irene Mendoza and Eva Moracho, for support in data collection and processing with some of the individual-based networks which was fundamental for this manuscript, as well as, discussion and comments for improving the quality of this work. We are thankful to Daniel Stouffer for his help and discussion on the network centralization metric, Giacomo Puglielli for encouraging ideas on PCA analysis and Enrico Bazzicalupo for his support and insights on cluster analysis. We thank the logistic and facilities support form ICTS-RBD Doñana and the Doñana National Park for the access authorizations to collect data from many of the individual networks here presented. We are thankful to two anonymous reviewers and the managing editor for their suggestions, which substantially helped to improve the manuscript. Funding from LIFEWATCH-2019-09-CSIC-13 project and PID2022-136812NB-I00, Spanish Ministry of Science and Innovation, as well as funding from the VI Plan Propio de Investigación of the Universidad de Sevilla (VI PPIT-US) supported this research (PJ). EQ was supported by “la Caixa’’ Foundation fellowship LCF/BQ/DE18/1167000. BAC received funding from the Ministry of Universities of the Spanish Government (Ref. FPU19_02552). JI was supported by FPI grant PRE2018-085916 from the Spanish Ministry of Science and Innovation. FRS was supported by VI PPIT-US and grants US-1381388 from Universidad de Sevilla/Junta de Andalucía/FEDER-UE and CNS2022-135839 funded by MICIU/AEI/10.13039/501100011033 and by European Union NextGenerationEU/PRTR.

