## Supplementary Information for "Downscaling mutualistic networks from species to individuals reveals consistent interaction niches and roles within plant populations"

[1] Integrative Ecology Group, Estación Biológica de Doñana, EBD-CSIC, Av. Américo Vespucio 26, E-41092 Sevilla, Spain.

[2] Departamento de Biología Vegetal y Ecología, Facultad de Biología, Universidad de Sevilla, Av. Reina Mercedes, E-41012 Sevilla, Spain.

### Network characteristics and sampling effort

**Table S1.** List of networks providing the identification code used in figures, the type (ind: individual-based, sp: species-based), the focal plant species in the case of individual-based networks, the country where network was sampled, number of plants and frugivore species present in the network, network size (potential interactions), number of unique interactions (realized interactions), the name of the population where sampling took place in case of several populations for the same study, and the reference of the study from which the network was extracted.

| Net no. | Type | Focal plant species | Country | Plants | Frugivores | Net size | Unique interactions | Population site | Sampling method | Reference |
|---|---|---|---|---|---|---|---|---|---|---|
| 1 | ind | *Pistacia lentiscus* (Anacardiaceae) | Spain | 40 | 27 | 1080 | 392 | El Puntal | Camera traps and DNA-barcoding | Quintero, E., Rodríguez-Sánchez, F., & Jordano, P. (2023). Reciprocity and interaction effectiveness in generalised mutualisms among free-living species. Ecology Letters, 26(1), 132–146. |
| 2 | ind | *Pistacia lentiscus* (Anacardiaceae) | Spain | 40 | 16 | 640 | 134 | Laguna de las Madroñas | DNA-barcoding | |
| 3 | ind | *Juniperus phoenicea* (Cupressaceae) | Spain | 35 | 10 | 350 | 137 | Colonizacion | Camera traps and DNA-barcoding | Isla, J., Jácome-Flores, M., Arroyo, J. M., & Jordano, P. (2023). The turnover of plant–frugivore interactions along plant range expansion: Consequences for natural colonization processes. Proceedings of the Royal Society B: Biological Sciences, 290(1999), 20222547. |
| 4 | ind | *Juniperus phoenicea* (Cupressaceae) | Spain | 35 | 10 | 350 | 154 | Ojillo | Camera traps and DNA-barcoding | |
| 5 | ind | *Juniperus phoenicea* (Cupressaceae) | Spain | 35 | 11 | 385 | 148 | El Marqués | Camera traps and DNA-barcoding | |
| 6 | ind | *Lithraea molleoides* (Anacardiaceae) | Argentina | 13 | 10 | 130 | 37 | Los Hornillos | Focal observations | Vergara-Tabares, D. L., Blendinger, P. G., Tello, A., Peluc, S. I., & Tecco, P. A. (2022). Fleshy-fruited invasive shrubs indirectly increase native tree seed dispersal. Oikos, 2022(2). |
| 7 | ind | *Lithraea molleoides* (Anacardiaceae) | Argentina | 14 | 10 | 140 | 33 | Las Calles | Focal observations | |
| 8 | ind | *Lithraea molleoides* (Anacardiaceae) | Argentina | 14 | 13 | 182 | 46 | La Poblacion | Focal observations | |
| 9 | ind | *Lithraea molleoides* (Anacardiaceae) | Argentina | 13 | 12 | 156 | 41 | San Javier | Focal observations | |

| Net no. | Type | Focal plant species | Country | Plants | Frugivores | Net size | Unique interactions | Population site | Sampling method | Reference |
|---|---|---|---|---|---|---|---|---|---|---|
| 10 | ind | *Lithraea molleoides* (Anacardiaceae) | Argentina | 12 | 11 | 132 | 29 | Las Rabonas | Focal observations | |
| 11 | ind | *Lithraea molleoides* (Anacardiaceae) | Argentina | 11 | 7 | 77 | 25 | Los Molles | Focal observations | |
| 12 | ind | *Laurus nobilis* (Lauraceae) | Spain | 18 | 17 | 306 | 87 | | Focal observations | Rodríguez-Sánchez, F. (2010). An integrative framework to investigate species responses to climate change: Biogeography and ecology of relict trees in the Mediterranean. PhD Thesis. Universidad de Sevilla, Spain. |
| 13 | ind | *Prunus mahaleb* (Rosaceae) | Spain | 19 | 20 | 380 | 211 | | Focal observations | Jordano, P. (1995). Frugivore-mediated selection on fruit and seed size: birds and St. Lucie's cherry, *Prunus mahaleb*. Ecology, 76(8), 2627–2639. Jordano, P., & Schupp, E. W. (2000). Seed disperser effectiveness: the quantity component and patterns of seed rain for *Prunus mahaleb*. Ecological Monographs, 70(4), 591-615. |
| 14 | ind | *Euterpe edulis* (Arecaceae) | Brazil | 17 | 9 | 153 | 31 | Restinga | Focal observations | Friedemann, P., Côrtes, M. C., de Castro, E. R., Galetti, M., Jordano, P., & Guimarães Jr, P. R. (2022). The individual-based network structure of palm-seed dispersers is explained by a rainforest gradient. Oikos, 2022, e08384. |
| 15 | ind | *Euterpe edulis* (Arecaceae) | Brazil | 15 | 7 | 105 | 25 | Lowland | Focal observations | |
| 16 | ind | *Euterpe edulis* (Arecaceae) | Brazil | 30 | 8 | 240 | 50 | Premontane | Focal observations | |
| 17 | ind | *Cecropia glaziovii* (Urticaceae) | Brazil | 27 | 37 | 999 | 124 | | Focal observations | Jordano, P. 2024. Material for the course: Curso Posgraduação Frugivoria 2016, UNESP Rio Claro, Brazil. doi: 10.5281/zenodo.10478535. |

| Net no. | Type | Focal plant species | Country | Plants | Frugivores | Net size | Unique interactions | Population site | Sampling method | Reference |
|---|---|---|---|---|---|---|---|---|---|---|
| 18 | ind | *Heynea trijuga* (Meliaceae) | India | 24 | 11 | 264 | 48 | | Focal observations | Gopal, A., Mudappa, D., Raman, T. S., & Naniwadekar, R. (2020). Forest cover and fruit crop size differentially influence frugivory of select rainforest tree species in Western Ghats, India. Biotropica, 52(5), 871-883. |
| 19 | ind | *Myristica dactyloides* (Myristicaceae) | India | 25 | 7 | 175 | 45 | | Focal observations | |
| 20 | ind | *Persea macrantha* (Lauraceae) | India | 32 | 21 | 672 | 186 | | Focal observations | |
| 21 | ind | *Henriettea succosa* (Melastomataceae ) | Brazil | 18 | 22 | 396 | 77 | | Focal observations | Crestani, A. C., Mello, M. A. R., & Cazetta, E. (2019). Interindividual variations in plant and fruit traits affect the structure of a plant-frugivore network. Acta Oecologica, 95, 120-127. |
| 22 | ind | *Prestoea decurrens* (Arecaceae) | Ecuador | 31 | 9 | 279 | 100 | | Camera traps | Lamperty, T., Karubian, J., & Dunham, A. E. (2021). Ecological drivers of intraspecific variation in seed dispersal services of a common neotropical palm. Biotropica, 53(4), 1226–1237. |
| 23 | ind | *Corema album* (Ericaceae) | Spain | 24 | 15 | 360 | 129 | | Camera traps | Villalva, P., Arroyo-Correa, B., Calvo, G., Homet, P., Isla, J., Mendoza, I., Moracho, E., Quintero, E., Rodríguez-Sánchez, F., & Jordano, P. (2023). FRUGIVORY CAMTRAP: A dataset of plant-animal interactions recorded with camera traps. https://doi.org/10.20350/DIGITALCSIC/15623 |
| 24 | ind | *Bursera penicillata* (Burseraceae) | India | 15 | 11 | 165 | 43 | | Focal observations | Ramaswami, G., Somnath, P., & Quader, S. (2017). Plant-disperser mutualisms in a semi-arid habitat invaded by Lantana camara L. Plant Ecology, 218, 935-946. |
| 25 | ind | *Erythroxylum monogynum* (Erythroxylaceae) | India | 12 | 6 | 72 | 29 | | Focal observations | |
| 26 | ind | *Flacourtia indica* (Salicaceae) | India | 13 | 5 | 65 | 33 | | Focal observations | |

| Net no. | Type | Focal plant species | Country | Plants | Frugivores | Net size | Unique interactions | Population site | Sampling method | Reference |
|---|---|---|---|---|---|---|---|---|---|---|
| 27 | ind | *Flueggea leucopyrus* (Phyllanthaceae) | India | 10 | 8 | 80 | 22 | | Focal observations | |
| 28 | ind | *Canthium coromandelicum* (Rubiaceae) | India | 10 | 8 | 80 | 30 | | Focal observations | |
| 29 | ind | *Santalum album* (Santalaceae) | India | 14 | 10 | 140 | 38 | | Focal observations | |
| 30 | ind | *Ziziphus oenopolia* (Rhamnaceae) | India | 15 | 13 | 195 | 102 | | Focal observations | |
| 31 | ind | *Chamaerops humilis* (Arecaceae) | Spain | 39 | 6 | 234 | 76 | Matasgordas | Footprint traps | Jácome-Flores, M. E. et al. 2020. Interaction motifs variability in a Mediterranean palm under environmental disturbances: the mutualism–antagonism continuum. Oikos 129: 367–379. |
| 32 | ind | *Chamaerops humilis* (Arecaceae) | Spain | 24 | 6 | 144 | 57 | Martinazo | Footprint traps | |
| 33 | ind | *Miconia irwinii* (Melastomataceae ) | Brazil | 15 | 9 | 135 | 59 | | Focal observations | Guerra, T. J. et al. 2017. Intraspecific variation in fruit–frugivore interactions: effects of fruiting neighborhood and consequences for seed dispersal. Oecologia 185: 233–243. |
| 34 | ind | *Juniperus macrocarpa* (Cupressaceae) | Spain | 26 | 11 | 286 | 72 | | Camera traps | Villalva, P., Arroyo-Correa, B., Calvo, G., Homet, P., Isla, J., Mendoza, I., Moracho, E., Quintero, E., Rodríguez-Sánchez, F., & Jordano, P. (2023). FRUGIVORY CAMTRAP: A dataset of plant-animal interactions recorded with camera traps. https://doi.org/10.20350/DIGITALCSIC/15623 |
| 35 | ind | *Prosopis flexuosa* (Fabaceae) | Argentina | 26 | 9 | 234 | 72 | Araya-San Ignacio | Camera traps | Miguel, M.F., Jordano, P., Tabeni, S. and Campos, C.M. (2018), Context-dependency and |

| Net no. | Type | Focal plant species | Country | Plants | Frugivores | Net size | Unique interactions | Population site | Sampling method | Reference |
|---|---|---|---|---|---|---|---|---|---|---|
| 36 | ind | *Prosopis flexuosa* (Fabaceae) | Argentina | 28 | 10 | 280 | 84 | El Bonito | Camera traps | anthropogenic effects on individual plant–frugivore networks. Oikos, 127: 1045-1059. |
| 37 | ind | *Prosopis flexuosa* (Fabaceae) | Argentina | 54 | 10 | 540 | 112 | El Doménico | Camera traps | |
| 38 | ind | *Prosopis flexuosa* (Fabaceae) | Argentina | 35 | 8 | 280 | 61 | MaB Ñacuñán Reserve Protected 1 | Camera traps | |
| 39 | ind | *Prosopis flexuosa* (Fabaceae) | Argentina | 29 | 9 | 261 | 99 | MaB Ñacuñán Reserve Protected 2 | Camera traps | |
| 40 | ind | *Schinus terebinthifolia* (Anacardiaceae) | Brazil | 26 | 16 | 416 | 93 | | Focal observations | Vissoto, M., Vizentin-Bugoni, J., Sendoya, S. F., Gomes, G. C., & Dias, R. A. (2022). Plant height and spatial context influence individual connectivity and specialization on seed dispersers in a tree population. Oecologia, 198(3): 721-731. |
| 41 | ind | *Phillyrea angustifolia* (Oleaceae) | Spain | 10 | 16 | 160 | 72 | El Puntal | DNA-barcoding | Moracho, E., Arroyo-Correa, B., Calvo, G., Homet, P., Isla, J., Mendoza, I., Quintero, E., Villalva, P., Rodríguez-Sánchez, F., & Jordano, P. (2023). DONANA-FRUGINT: A dataset of plant-animal interactions. https://doi.org/10.20350/DIGITALCSIC/#####. |
| 42 | ind | *Phillyrea angustifolia* (Oleaceae) | Spain | 9 | 12 | 108 | 41 | Laguna de las Madroñas | DNA-barcoding | |
| 43 | ind | *Marcgravia longifolia* (Marcgraviaceae) | Peru | 24 | 43 | 1032 | 127 | | Focal observations | Thiel, S., Willems, F., Farwig, N., Rehling, F., Schabo, D. G., Schleuning, M., Shahuano Tello, N., Töpfer, T., Tschapka, M., Heymann, E. W., & Heer, K. (2023). Vertically stratified frugivore community composition and interaction frequency in a liana fruiting across forest strata. Biotropica, 55, 650–664 |

| Net no. | Type | Focal plant species | Country | Plants | Frugivores | Net size | Unique interactions | Population site | Sampling method | Reference |
|---|---|---|---|---|---|---|---|---|---|---|
| 44 | ind | *Osyris lanceolata* (Santalaceae) | Spain | 19 | 14 | 266 | 62 | | DNA-barcoding | Moracho, E., Arroyo-Correa, B., Calvo, G., Homet, P., Isla, J., Mendoza, I., Quintero, E., Villalva, P., Rodríguez-Sánchez, F., & Jordano, P. (2023). DONANA-FRUGINT: A dataset of plant-animal interactions. https://doi.org/10.20350/DIGITALCSIC/#####. |
| 45 | ind | *Naringi crenulata* (Rutaceae) | India | 22 | 12 | 264 | 62 | | Focal observations | Jayanth, A., Isvaran, K., & Naniwadekar, R. (2024). Drivers of intraspecific variation in seed dispersal can differ across two species of fleshy-fruited savanna plants. Biotropica, 56(3), e13322. https://doi.org/10.1111/btp.13322 |
| 46 | ind | *Ziziphus oenopolia* (Rhamnaceae) | India | 20 | 13 | 260 | 57 | | Focal observations and mist-netting | |
| 47 | sp | | Spain | 25 | 36 | 900 | 228 | Cazorla | Focal observations | Olesen JM, Bascompte J, Dupont YL, Elberling H, Rasmussen C, Jordano P. (2011). Missing and forbidden links in mutualistic networks. Proceedings of the Royal Society B-Biological Sciences 278: 725-732. |
| 48 | sp | | Spain | 16 | 17 | 272 | 120 | Hato Ratón | Mist-netting | |
| 49 | sp | | Spain | 17 | 28 | 476 | 130 | Spain | Focal observations | García-Castaño, J.L. (2001). Consecuencias demográficas de la dispersión de semillas por aves y mamíferos frugívoros en la vegetación mediterránea de montaña. PhD Thesis. Universidad de Sevilla, Spain. |
| 50 | sp | | Papua New Guinea | 31 | 9 | 279 | 119 | | Focal observations | Beehler B. (1983) Frugivory and polygamy in birds of paradise. Auk, 100, 1-11. |
| 51 | sp | | South Africa | 16 | 10 | 160 | 110 | | Focal observations | Frost P.G.H. (1980) Fruit-frugivore interactions in a South African coastal dune forest. In: Acta XVII Congressus Internationalis Ornithologici (ed. Noring R), pp. 1179-1184. Deutsche Ornithologische Ges., Berlin, Germany. |

| Net no. | Type | Focal plant species | Country | Plants | Frugivores | Net size | Unique interactions | Population site | Sampling method | Reference |
|---|---|---|---|---|---|---|---|---|---|---|
| 52 | sp | | Spain | 12 | 7 | 84 | 40 | | Mist-netting | Guitián J. (1983) Relaciones entre los frutos y los passeriformes en un bosque montano de la cordillera Cantabrica occidental. PhD Thesis. Universidad de Santiago, Spain. |
| 53 | sp | | Brazil | 35 | 29 | 1015 | 146 | | Focal observations | Galetti M. & Pizo M.A. (1996) Fruit eating birds in a forest fragment in southeastern Brazil. Ararajuba, Rev. Brasil. Ornitol., 4, 71-79. |
| 54 | sp | | England | 11 | 14 | 154 | 47 | | Focal observations | Snow, B.K. & Snow, D.W. (1988). Birds and berries. T. and A.D. Poyser, Calton, England. |
| 55 | sp | | Japan | 15 | 8 | 120 | 38 | | Focal observations | Noma, N. & Yumoto, T. (1997). Fruiting phenology of animal-dispersed plants in response to winter migration of frugivores in a warm temperate forest on Yakushima Island, Japan. Ecological Research, 12, 119-129. |
| 56 | sp | | Australia | 71 | 7 | 497 | 142 | | Focal observations | Crome, F. H.J. (1975). The ecology of fruit pigeons in tropical Northern Queensland. Aust Wildl Res, 2, 155-185. |
| 57 | sp | | Trinidad and Tobago | 50 | 14 | 700 | 234 | | Focal observations | Snow B.K. & Snow D.W. (1971) The feeding ecology of tanagers and honeycreepers in Trinidad. Auk, 88, 291-322. |
| 58 | sp | | United States | 7 | 21 | 147 | 50 | | Focal observations | Baird, J.W. (1980). The selection and use of fruit by birds in an Eastern forest. Wilson Bulletin, 92, 63-73. |
| 59 | sp | | Kenya | 8 | 30 | 240 | 69 | Interior little disturbed | Focal observations | Menke, S., Böhning-Gaese, K. & Schleuning, M. (2012). Plant-frugivore networks are less specialized and more robust at forest-farmland edges than in the interior of a tropical forest. Oikos, 121, 1553-1566. |
| 60 | sp | | Kenya | 7 | 38 | 266 | 104 | Edge little disturbed | Focal observations | |
| 61 | sp | | Kenya | 8 | 34 | 272 | 88 | Interior highly disturbed | Focal observations | |

| Net no. | Type | Focal plant species | Country | Plants | Frugivores | Net size | Unique interactions | Population site | Sampling method | Reference |
|---|---|---|---|---|---|---|---|---|---|---|
| 62 | sp | | Kenya | 8 | 39 | 312 | 115 | Edge highly disturbed | Focal observations | |
| 63 | sp | | Brazil | 15 | 49 | 735 | 143 | | Focal observations | Pizo, M.A. (2004). Frugivory and habitat use by fruit-eating birds in a fragmented landscape of southeast Brazil. Ornitologia Neotropical, 15, 117-126. |
| 64 | sp | | Kenya | 33 | 88 | 2904 | 419 | | Focal observations | Schleuning M, Bluthgen N, Florchinger M, Braun J, Schaefer HM, Bohning-Gaese K. 2011. Specialization and interaction strength in a tropical plant-frugivore network differ among forest strata. Ecology 92: 26-36. |
| 65 | sp | | Brazil | 49 | 16 | 784 | 131 | | Focal observations | Castro, E.R.D. (2007). Fenologia reprodutiva do palmito Euterpe edulis (Erecaceae) e sua influência na abundância de aves frugívoras na floresta atlântica. PhD Thesis. Instituto de Biociencias. Universidade Estadual Paulista "Juio de Mesquita Filho" Rio Claro, SP, Brazil. |
| 66 | sp | | Brazil | 13 | 45 | 585 | 183 | | Focal observations | Correia, J.M.S. (1997). Utilização de espécies frutíferas de mata Atlântica na alimentação da avifauna da reserva biológica de Poço das Antas, RJ. MSc Thesis. Instituto de Biologia. UNB, Brazil. |
| 67 | sp | | Brazil | 13 | 30 | 390 | 145 | | Focal observations | Alves, K.J.F. (2008). Composição da avifauna e frugivoria por aves em um mosaico sucessional na mata Atlântica. MSc Thesis. Instituto de Biociencias. Universidade Estadual Paulista, Julio de Mesquita Filho, Rio Claro, SP, Brazil. |
| 68 | sp | | Brazil | 9 | 30 | 270 | 92 | | Focal observations | Athie, S. (2009). Composição da avifauna e frugivoria por aves em um mosaico de vegetação secundária em Rio Claro, região centro-leste do estado de São Paulo. MSc Thesis. Universidade Federal de São Carlos, Brazil. |

| Net no. | Type | Focal plant species | Country | Plants | Frugivores | Net size | Unique interactions | Population site | Sampling method | Reference |
|---|---|---|---|---|---|---|---|---|---|---|
| 69 | sp | | Brazil | 25 | 28 | 700 | 90 | | Focal observations | Ferreira Fadini, R. & De Marco Jr., P. (2004). Interações entre aves frugívoras e plantas em um fragmento de mata atlântica de Minas Gerais. Ararajuba, 12, 97-103. |
| 70 | sp | | Brazil | 26 | 22 | 572 | 79 | | Focal observations and mist-netting | Hasui, Erica. (1994). O papel das aves frugivoras na dispersao de sementes em um fragmento de floresta semidecidua secundaria em Sao Paulo, SP. MSc thesis. USP Sao Paulo, Brazil. |
| 71 | sp | | Brazil | 22 | 20 | 440 | 67 | | Focal observations | Silva, R. F. d. M. (2011). Interações entre plantas e aves frugívoras no campus da Universidade Federal Rural do Rio de Janeiro. In: Instituto de Florestas. Universidade Federal do Rio de Janeiro Rio de Janeiro, Brazil. |
| 72 | sp | | Brazil | 12 | 15 | 180 | 32 | 15-year-old restored plot | Focal observations | Ribeiro da Silva, F., Montoya, D., Furtado, R., Memmott, J., Pizo, M.A. and Rodrigues, R.R. (2015), The restoration of tropical seed dispersal networks. Restor Ecol, 23: 852-860. |
| 73 | sp | | Brazil | 23 | 29 | 667 | 129 | 25-year-old restored plot | Focal observations | |
| 74 | sp | | Brazil | 14 | 14 | 196 | 35 | 57-year-old restored plot | Focal observations | |
| 75 | sp | | Brazil | 6 | 28 | 168 | 50 | | Focal observations | Robinson, V. (2015). Interações entre aves frugívoras e plantas em um fragmento de mata atlântica de Minas Gerais. In: Instituto de Biociencias. Universidade Estadual Paulista "Julio de Mesquita Filho" Rio Claro, SP, Brazil. |
| 76 | sp | | Brazil | 30 | 58 | 1740 | 240 | | Focal observations | Rodrigues, S. B. M. (2015). Rede de interações entre aves frugivoras e plantas em una área de mata Atlântica no sudeste do Brasil. Universidade Federal de Sao Carlos, Campus Sorocaba Sorocaba, SP, Brazil. |

| Net no. | Type | Focal plant species | Country | Plants | Frugivores | Net size | Unique interactions | Population site | Sampling method | Reference |
|---|---|---|---|---|---|---|---|---|---|---|
| 77 | sp | | New Zealand | 18 | 8 | 144 | 102 | | Focal observations | Burns, K.C. (2013). What causes size coupling in fruit-frugivore interaction webs? Ecology, 94, 295-300. |
| 78 | sp | | Poland | 8 | 12 | 96 | 38 | site_11 | Focal observations | Albrecht, J., Bohle, V., Berens, D. G., Jaroszewicz, B., Selva, N., & Farwig, N. (2015). Variation in neighbourhood context shapes frugivore-mediated facilitation and competition among co-dispersed plant species. Journal of Ecology, 103(2), 526–536. |
| 79 | sp | | Poland | 7 | 11 | 77 | 32 | site_13 | Focal observations | |
| 80 | sp | | Poland | 9 | 13 | 117 | 42 | site_15 | Focal observations | |
| 81 | sp | | Poland | 8 | 13 | 104 | 36 | site_30 | Focal observations | |
| 82 | sp | | Poland | 8 | 10 | 80 | 29 | site_35 | Focal observations | |
| 83 | sp | | Poland | 8 | 10 | 80 | 30 | site_36 | Focal observations | |
| 84 | sp | | Poland | 8 | 11 | 88 | 33 | site_102 | Focal observations | |
| 85 | sp | | Poland | 10 | 13 | 130 | 42 | site_111 | Focal observations | |
| 86 | sp | | Poland | 8 | 15 | 120 | 57 | site_112 | Focal observations | |
| 87 | sp | | Poland | 9 | 16 | 144 | 41 | site_203 | Focal observations | |
| 88 | sp | | Poland | 6 | 20 | 120 | 43 | site_301 | Focal observations | |
| 89 | sp | | Poland | 8 | 19 | 152 | 56 | site_315 | Focal observations | |
| 90 | sp | | Brazil | 22 | 17 | 374 | 78 | | Focal observations | Andrade, P., Mota, J. & Carvalho, A. (2011). Mutual interactions between frugivorous birds and plants in an urban fragment of Atlantic Forest, Salvador, BA. Revista Brasileira de Ornitologia, 19, 63-73. |

| Net no. | Type | Focal plant species | Country | Plants | Frugivores | Net size | Unique interactions | Population site | Sampling method | Reference |
|---|---|---|---|---|---|---|---|---|---|---|
| 91 | sp | | Puerto Rico | 34 | 20 | 680 | 95 | | Focal observations | Yang, S., Albert, R. & Carlo, T.A. (2013). Transience and constancy of interactions in a plant-frugivore network. Ecosphere, 4(12): 147. |
| 92 | sp | | Brazil | 14 | 6 | 84 | 22 | | Mist-netting | Garcia, Q.S., Rezende, J.L.P. & Aguiar, L.M.S. (2000). Seed dispersal by bats in a disturbed area of Southeastern Brazil. Revista de Biología Tropical, 48, 125-128. |
| 93 | sp | | Peru | 77 | 18 | 1386 | 196 | | Mist-netting | Gorchov, D.L., Cornejo, F., Ascorra, C.F. & Jaramillo, M. (1995). Dietary overlap between frugivorous birds and bats in the peruvian amazon. Oikos, 74, 235-250. |
| 94 | sp | | Costa Rica | 35 | 14 | 490 | 95 | | Mist-netting | Palmeirim, J.M., Gorchov, D.L. & Stoleson, S. (1989). Trophic structure of a neotropical frugivore community: is there competition between birds and bats? Oecologia (Berl.), 79, 403-411. |
| 95 | sp | | Costa Rica | 35 | 14 | 490 | 119 | | Mist-netting | Lopez, J.E. & Vaughan, C. (2004). Observations on the Role of Frugivorous Bats as Seed Dispersers in Costa Rican Secondary Humid Forests. Acta Chiropterologica, 6, 111-119. |
| 96 | sp | | Ecuador | 43 | 15 | 645 | 97 | | Animal feces (mist-netting and transects) | Heleno, R.H., Olesen, J.M., Nogales, M., Vargas, P. & Traveset, A. (2013). Seed dispersal networks in the Galapagos and the consequences of alien plant invasions. Proc Biol Sci, 280, 20122112. |
| 97 | sp | | Mexico | 22 | 7 | 154 | 47 | Tropical montane cloud forest fragment | Mist-netting | Hernandez-Montero, J.R., Saldana-Vazquez, R.A., Galindo-Gonzalez, J. & Sosa, V.J. (2015). Bat-fruit interactions are more specialized in shaded-coffee plantations than in tropical mountain cloud forest fragments. PLoS ONE, 10, e0126084. |
| 98 | sp | | Mexico | 19 | 6 | 114 | 34 | Shaded-coffee plantation | Mist-netting | |

| Net no. | Type | Focal plant species | Country | Plants | Frugivores | Net size | Unique interactions | Population site | Sampling method | Reference |
|---|---|---|---|---|---|---|---|---|---|---|
| 99 | sp | | Brazil | 24 | 7 | 168 | 50 | | Mist-netting | Passos, F.C., Silva, W.R., Pedro, W.A. & Bonin, M.R. (2003). Frugivoria em morcegos (Mammalia, Chiroptera) no Parque Estadual Intervales, sudeste do Brasil. Revista Brasileira de Zoologia, 20, 511-517. |
| 100 | sp | | Brazil | 13 | 7 | 91 | 30 | | Mist-netting | Pedro, W.A. (1992). Estrutura de uma taxocenose de morcegos da reserva do Panga (Uberlandia, MG), com enfase nas relações troficas em Phyllostomidae (Mammalia: Chiroptera). MSC thesis. Universidade Estadual de Campinas Campinas, SP, Brazil. |
| 101 | sp | | Panama | 17 | 20 | 340 | 86 | | Mist-netting | Poulin, B., Wright, S.J., Lefebvre, G. & Calderon, O. (1999). Interspecific synchrony and asynchrony in the fruiting phenologies of congeneric bird-dispersed plants in Panama. Journal of Tropical Ecology, 15, 213-227. |
| 102 | sp | | Brazil | 56 | 20 | 1120 | 104 | | Mist-netting | Sarmento, R., Alves-Costa, C.i.P., Ayub, A. & Mello, M.A.R. (2014). Partitioning of seed dispersal services between birds and bats in a fragment of the Brazilian Atlantic Forest. Zoologia (Curitiba, Impresso), 31, 245-255. |
| 103 | sp | | Germany | 30 | 31 | 930 | 189 | | Focal observations | Stiebel, H. & Bairlein, F. (2008). Frugivorie mitteleuropäischer Vögel I: Nahrung und Nahrungserwerb. Vogelwarte, 46, 1-23. |
| 104 | sp | | Bolivia | 36 | 41 | 1476 | 127 | Forest edge | Focal observations | Saavedra, F., Hensen, I., Beck, S. G., Böhning-Gaese, K., Lippok, D., Töpfer, T., & Schleuning, M. (2014). Functional importance of avian seed dispersers changes in response to human-induced forest edges in tropical seed-dispersal networks. Oecologia, 176, 837-848. |
| 105 | sp | | Bolivia | 20 | 23 | 460 | 52 | Forest interior | Focal observations | |

**Table S2.** Sampling coverage in individual-based networks with lower- and upper-confidence limits of sample coverage (95%) (SC LCL, SC UCL, respectively).

| Net no. | Focal plant species | Sampling coverage | SC LCL | SC UCL |
|---|---|---|---|---|
| 1 | *Pistacia lentiscus* | 0.99 | 0.99 | 1.00 |
| 2 | *Pistacia lentiscus* | 0.96 | 0.93 | 0.99 |
| 3 | *Juniperus phoenicea* | 0.99 | 0.97 | 1.00 |
| 4 | *Juniperus phoenicea* | 0.99 | 0.98 | 1.00 |
| 5 | *Juniperus phoenicea* | 1.00 | 0.99 | 1.00 |
| 6 | *Lithraea molleoides* | 0.87 | 0.78 | 0.96 |
| 7 | *Lithraea molleoides* | 0.94 | 0.85 | 1.00 |
| 8 | *Lithraea molleoides* | 0.92 | 0.86 | 0.98 |
| 9 | *Lithraea molleoides* | 0.91 | 0.82 | 1.00 |
| 10 | *Lithraea molleoides* | 0.87 | 0.74 | 1.00 |
| 11 | *Lithraea molleoides* | 0.93 | 0.82 | 1.00 |
| 12 | *Laurus nobilis* | 0.98 | 0.95 | 1.00 |
| 13 | *Prunus mahaleb* | 1.00 | 0.99 | 1.00 |
| 14 | *Euterpe edulis* | 0.91 | 0.80 | 1.00 |
| 15 | *Euterpe edulis* | 0.89 | 0.79 | 0.99 |
| 16 | *Euterpe edulis* | 0.96 | 0.92 | 1.00 |
| 17 | *Cecropia glaziovii* | 0.88 | 0.84 | 0.93 |
| 18 | *Heynea trijuga* | 0.96 | 0.91 | 1.00 |
| 19 | *Myristica dactyloides* | 0.98 | 0.94 | 1.00 |
| 20 | *Persea macrantha* | 0.98 | 0.96 | 1.00 |
| 21 | *Henriettea succosa* | 0.87 | 0.81 | 0.93 |
| 22 | *Prestoea decurrens* | 0.98 | 0.97 | 1.00 |
| 23 | *Corema album* | 0.98 | 0.95 | 1.00 |
| 24 | *Bursera penicillata* | 0.94 | 0.87 | 1.00 |
| 25 | *Erythroxylum monogynum* | 0.97 | 0.91 | 1.00 |
| 26 | *Flacourtia indica* | 1.00 | 0.97 | 1.00 |
| 27 | *Flueggea leucopyrus* | 0.84 | 0.72 | 0.95 |
| 28 | *Canthium coromandelicum* | 0.91 | 0.82 | 1.00 |
| 29 | *Santalum album* | 0.88 | 0.79 | 0.96 |
| 30 | *Ziziphus oenopolia* | 0.97 | 0.94 | 1.00 |
| 31 | *Chamaerops humilis* | 1.00 | 0.99 | 1.00 |
| 32 | *Chamaerops humilis* | 1.00 | 0.99 | 1.00 |
| 33 | *Miconia irwinii* | 1.00 | 0.98 | 1.00 |
| 34 | *Juniperus macrocarpa* | 0.97 | 0.94 | 1.00 |
| 35 | *Prosopis flexuosa* | 1.00 | 0.98 | 1.00 |
| 36 | *Prosopis flexuosa* | 0.97 | 0.93 | 1.00 |
| 37 | *Prosopis flexuosa* | 0.97 | 0.95 | 0.99 |
| 38 | *Prosopis flexuosa* | 0.95 | 0.92 | 0.99 |
| 39 | *Prosopis flexuosa* | 0.99 | 0.97 | 1.00 |
| 40 | *Schinus terebinthifolia* | 0.96 | 0.92 | 0.99 |
| 41 | *Phillyrea angustifolia* | 0.95 | 0.90 | 0.99 |
| 42 | *Phillyrea angustifolia* | 0.94 | 0.88 | 1.00 |
| 43 | *Marcgravia longifolia* | 0.82 | 0.75 | 0.88 |
| 44 | *Osyris lanceolata* | 0.90 | 0.84 | 0.97 |
| 45 | *Naringi crenulata* | 0.96 | 0.90 | 1.00 |
| 46 | *Ziziphus oenopolia* | 0.93 | 0.87 | 1.00 |

### Comparison of networks at different resolution scales

To compare networks focused at the population level (individual-based) and the community level (species-based) we calculated several network descriptors. We then used these descriptors to build a PCA-derived multivariate space defined by their correlation structure, so that the location of each network is defined by a combination of both topological (e.g., degree, connectance) and structural (e.g., nestedness, modularity) descriptors. In this way, networks closely located in this multivariate space would have more similarities in the combination of metrics (and thus topology and structure) than networks located in different parts of the space.

#### *Network-level metrics*

With the aim of visualizing families of metrics that describe similar aspects of the bipartite networks, we computed an agglomerative hierarchical clustering (HC) analysis for all the metrics (function hclust in `stats` R package, R Core Team 2023).

We selected metrics indicative of biological properties of the networks, aiming to reduce redundancy in their meaning and avoiding high correlation with network size. Since we aim at finding structural differences among networks with different resolution scales we tried to avoid metrics strongly affected by sampling design, species diversity and study region characteristics (e.g., tropical vs. temperate regions), such as web asymmetry, Shannon diversity or links per species. Both the cluster analysis and the correlation analysis help us select network-level metrics that are interpretable in biological terms while trying to avoid highly correlated metrics. The selected network-level metrics allow us to discern differences in the topological properties of individual-based and species-based networks.

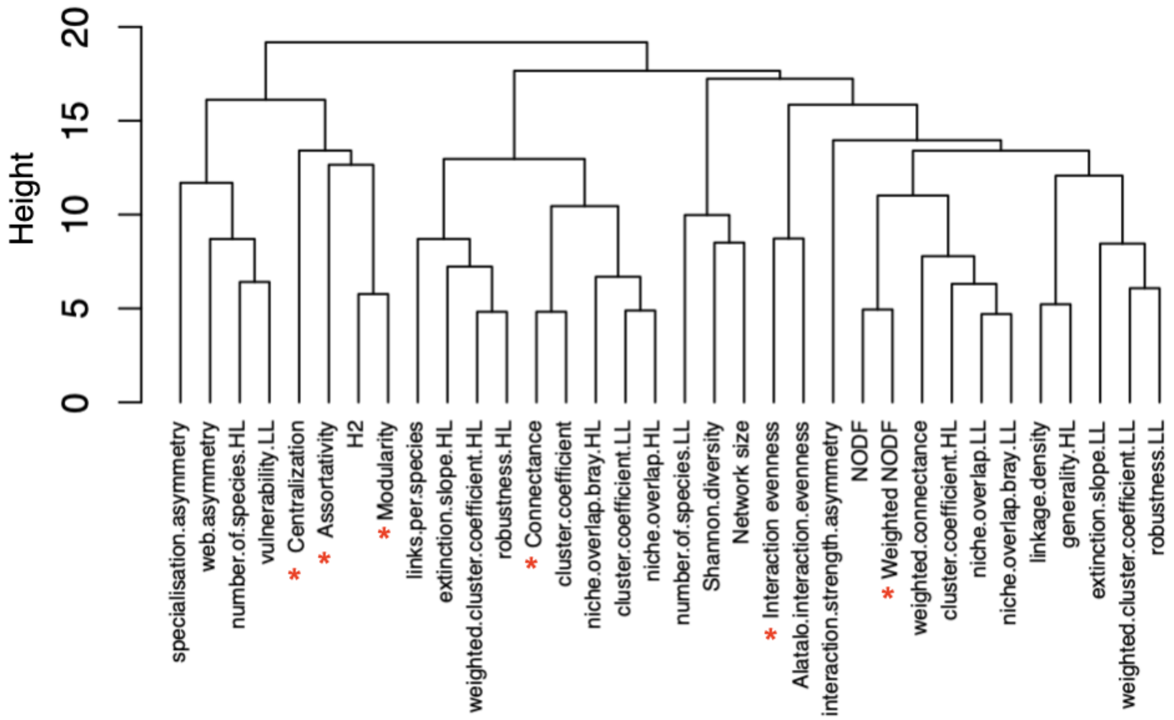


**Figure S1.** Hierarchical clustering analysis results for all the network metrics calculated. Metrics with * are the selected ones.

We checked for Pearson's correlation among the selected metrics and with network size (Fig. S2). We did not find strong effects of correlation with network size (medium/low correlation). The highest correlations were between centralization and interaction evenness, and weighted NODF and modularity. All variables have a VIF of 2.52 (VIF < 3).

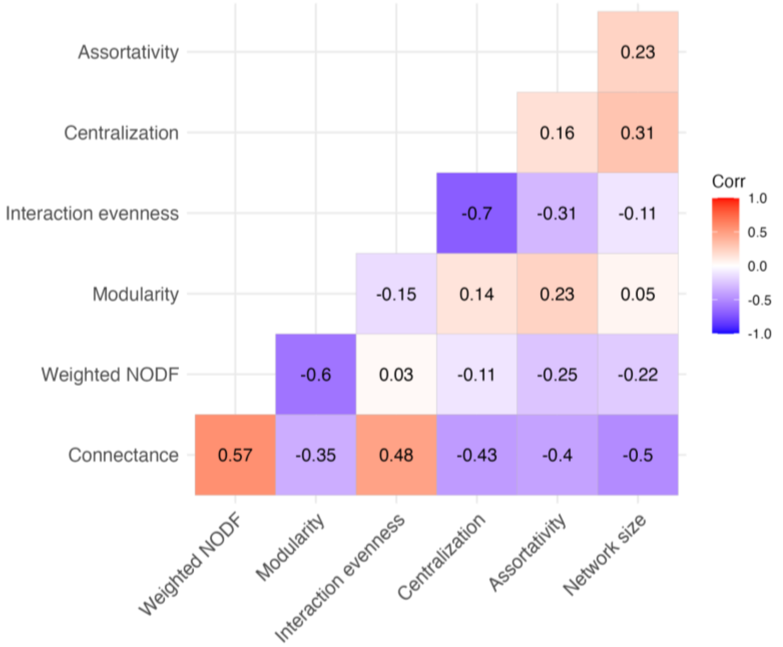


**Figure S2.** Correlation plot between selected network-level metrics for PCA analysis. Numbers denote Pearson correlation coefficients (r) and the color its magnitude and direction.

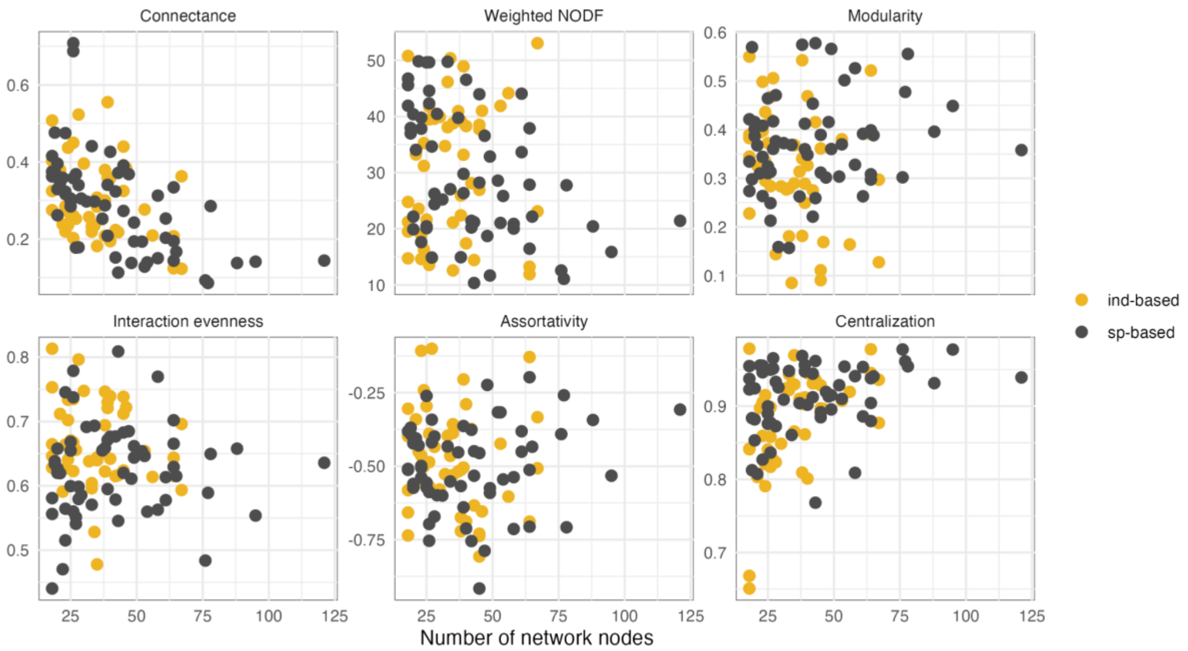


**Figure S3.** Correlation between selected network metrics and the number of nodes present in the network (species and/or individuals).

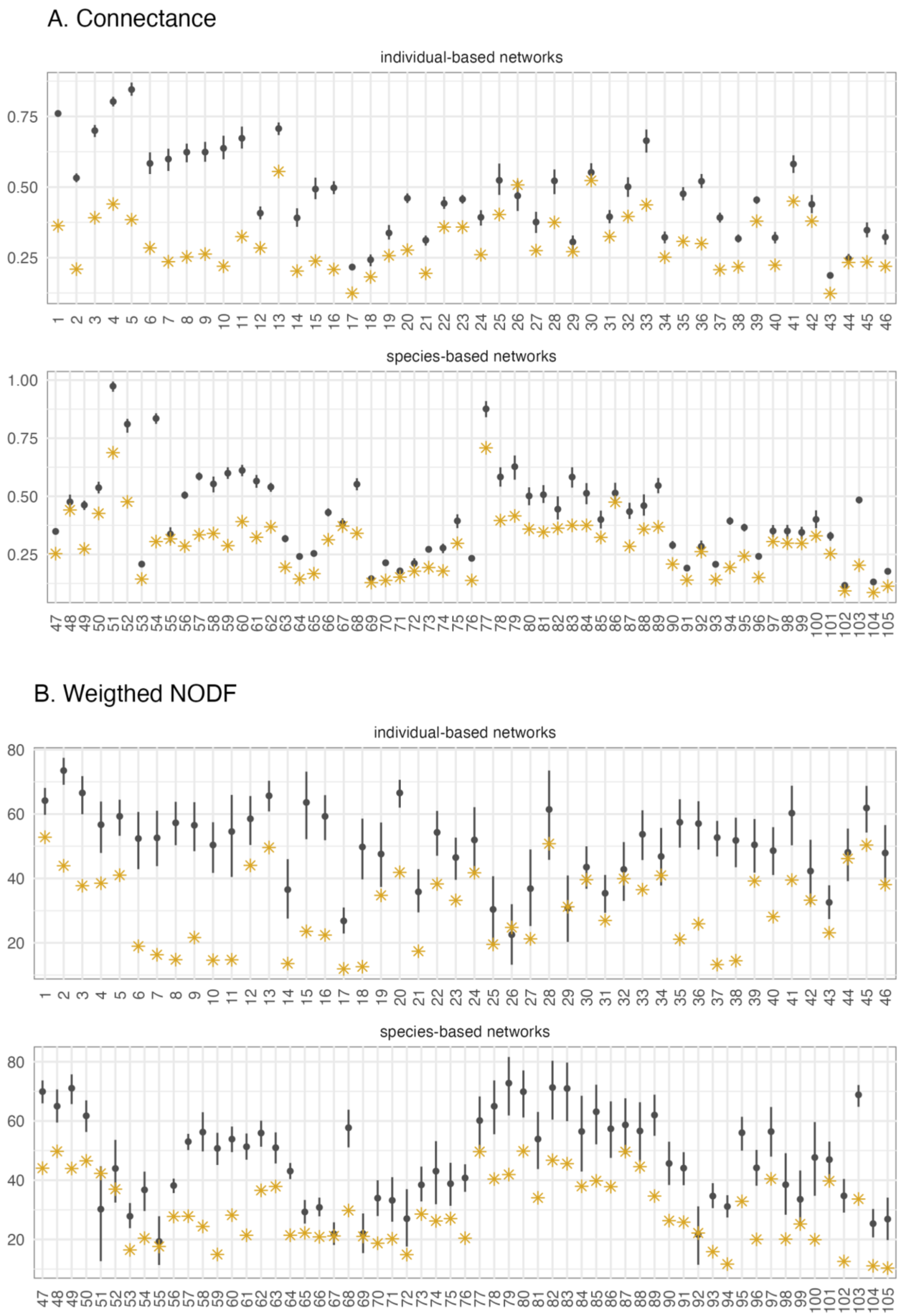
A. Connectance
individual-based networks
0.75
0.50
0.25
species-based networks
1.00
0.75
0.50
0.25
B. Weigthed NODF
individual-based networks
80
60
40
20
species-based networks
80
60
40
20

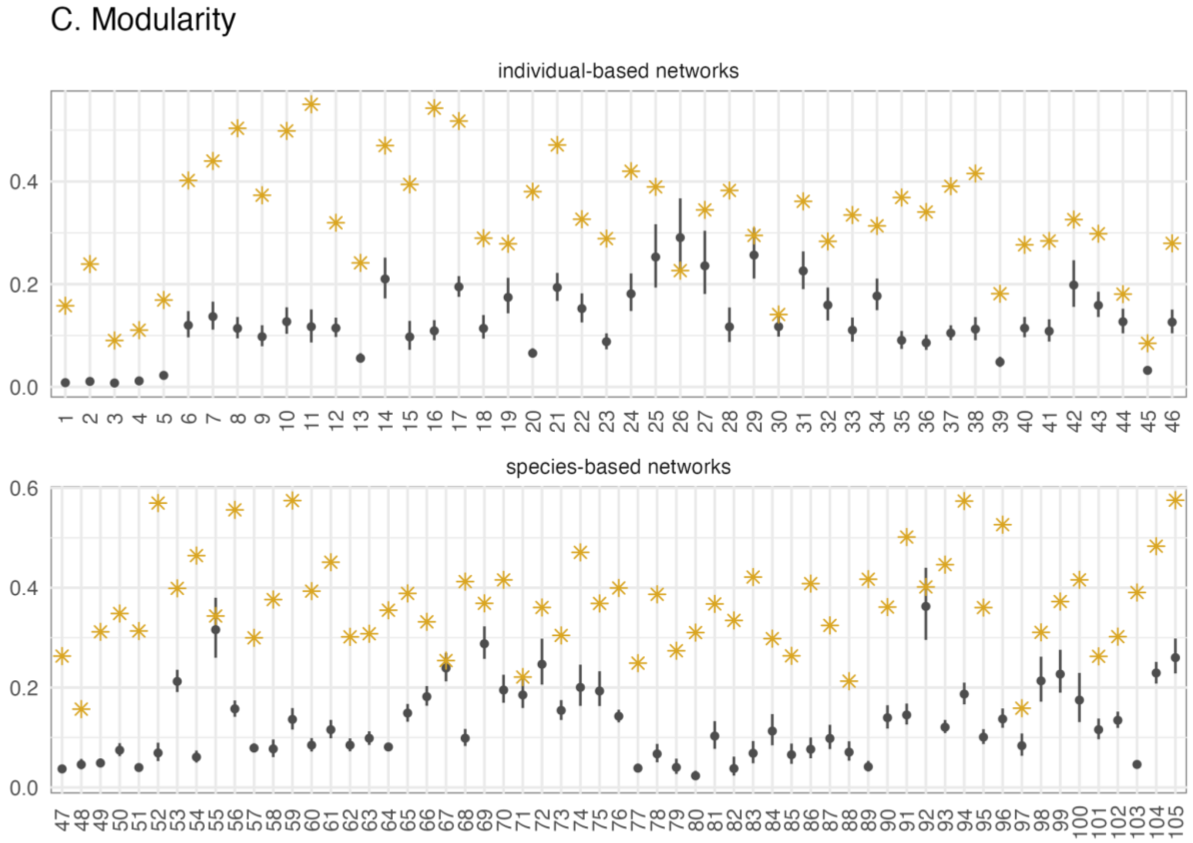
C. Modularity
individual-based networks
species-based networks

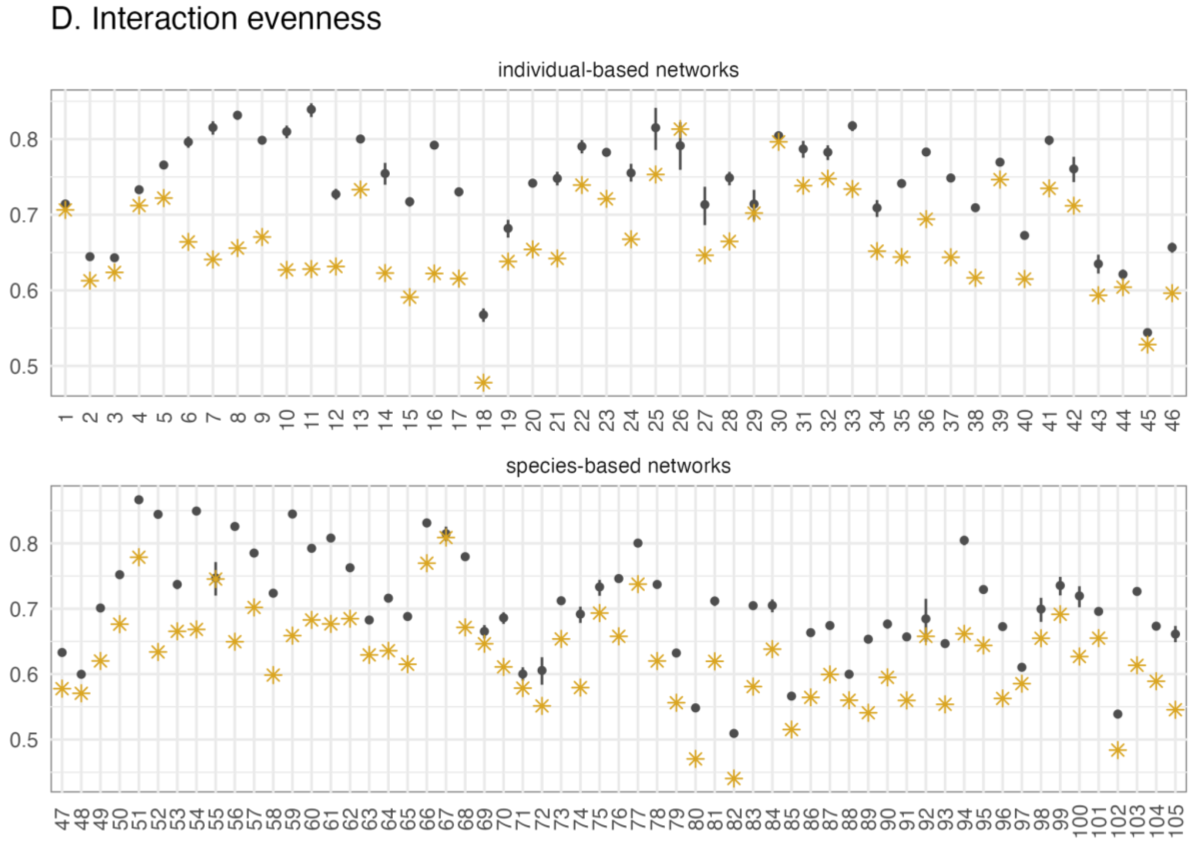
D. Interaction evenness
individual-based networks
species-based networks

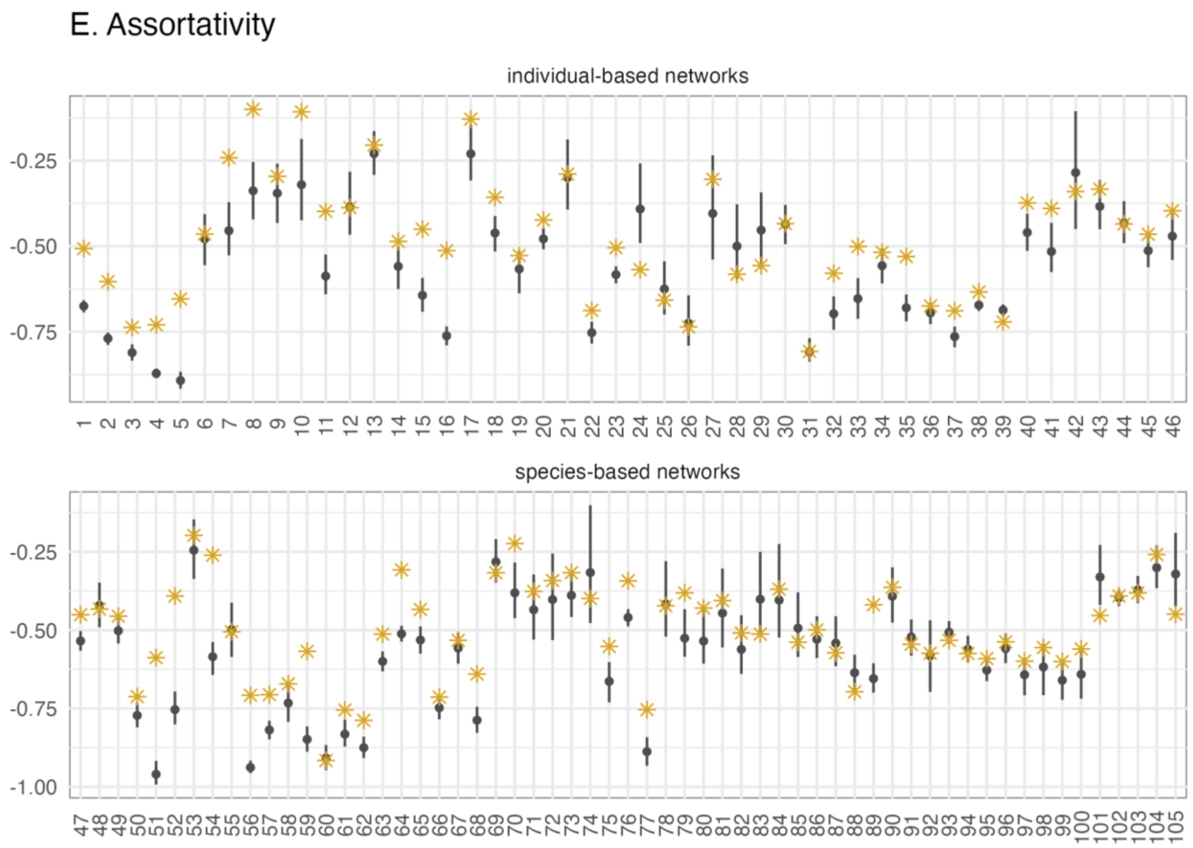
E. Assortativity
individual-based networks
-0.25
-0.50
-0.75
species-based networks
-0.25
-0.50
-0.75
-1.00

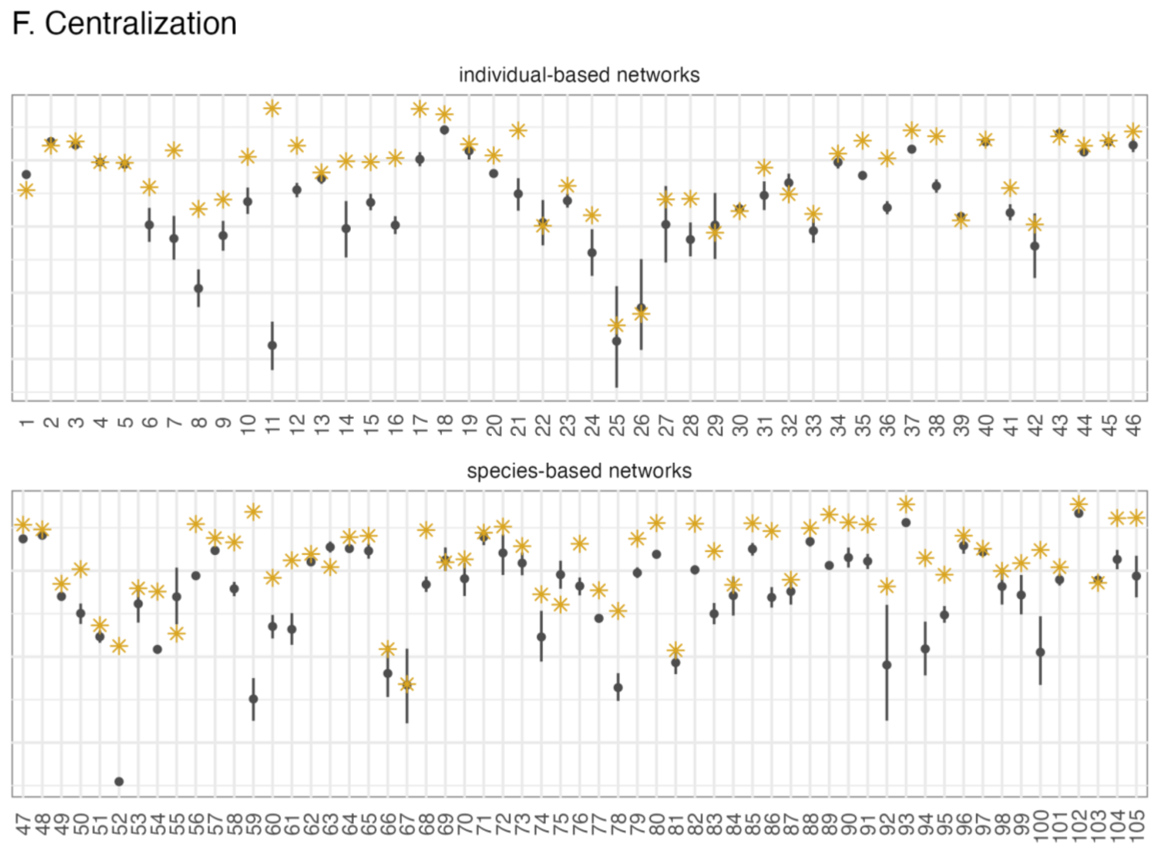
F. Centralization
individual-based networks
0.9
0.8
0.7
0.6
species-based networks
0.9
0.8
0.7

**Figure S4.** Network-level metrics for individual-based and species-based networks. Yellow asterisks represent empirical values and gray points intervals represent null model values (mean and 95% CI). Networks whose empirical value falls outside the confidence intervals are significantly different to their random expectation ($p > 0.05$). The null model algorithm used is Patefield algorithm which preserves marginal totals.

**Table S3.** Average metric values and standard deviation (SD) for species-based (sp) and individual-based (ind) networks. Null networks were generated using Patefield's null model algorithm that maintains network size (number of rows and columns) and interaction abundances (marginal totals).

| Metric | Type | Observed mean ± SD | Nulls mean ± SD | Average difference ± SD |
|---|---|---|---|---|
| Connectance | ind | 0.3 ± 0.1 | 0.48 ± 0.01 | -0.18 ± 0.12 |
| | sp | 0.29 ± 0.13 | 0.42 ± 0.01 | -0.13 ± 0.1 |
| Weighted NODF | ind | 30.52 ± 12.26 | 50.48 ± 4.05 | -19.96 ± 12.68 |
| | sp | 29.66 ± 11.53 | 46.96 ± 3.61 | -17.29 ± 9.32 |
| Modularity | ind | 0.33 ± 0.12 | 0.13 ± 0.01 | 0.2 ± 0.11 |
| | sp | 0.37 ± 0.1 | 0.13 ± 0.01 | 0.24 ± 0.11 |
| Interaction evenness | ind | 0.67 ± 0.07 | 0.74 ± 0 | -0.08 ± 0.05 |
| | sp | 0.62 ± 0.07 | 0.71 ± 0 | -0.08 ± 0.04 |
| Assortativity | ind | -0.48 ± 0.17 | -0.55 ± 0.03 | 0.07 ± 0.09 |
| | sp | -0.5 ± 0.15 | -0.57 ± 0.04 | 0.06 ± 0.11 |
| Centralization | ind | 0.88 ± 0.07 | 0.84 ± 0.01 | 0.04 ± 0.06 |
| | sp | 0.91 ± 0.05 | 0.87 ± 0.01 | 0.04 ± 0.04 |

### Population specialization (TNW ~ WIC)

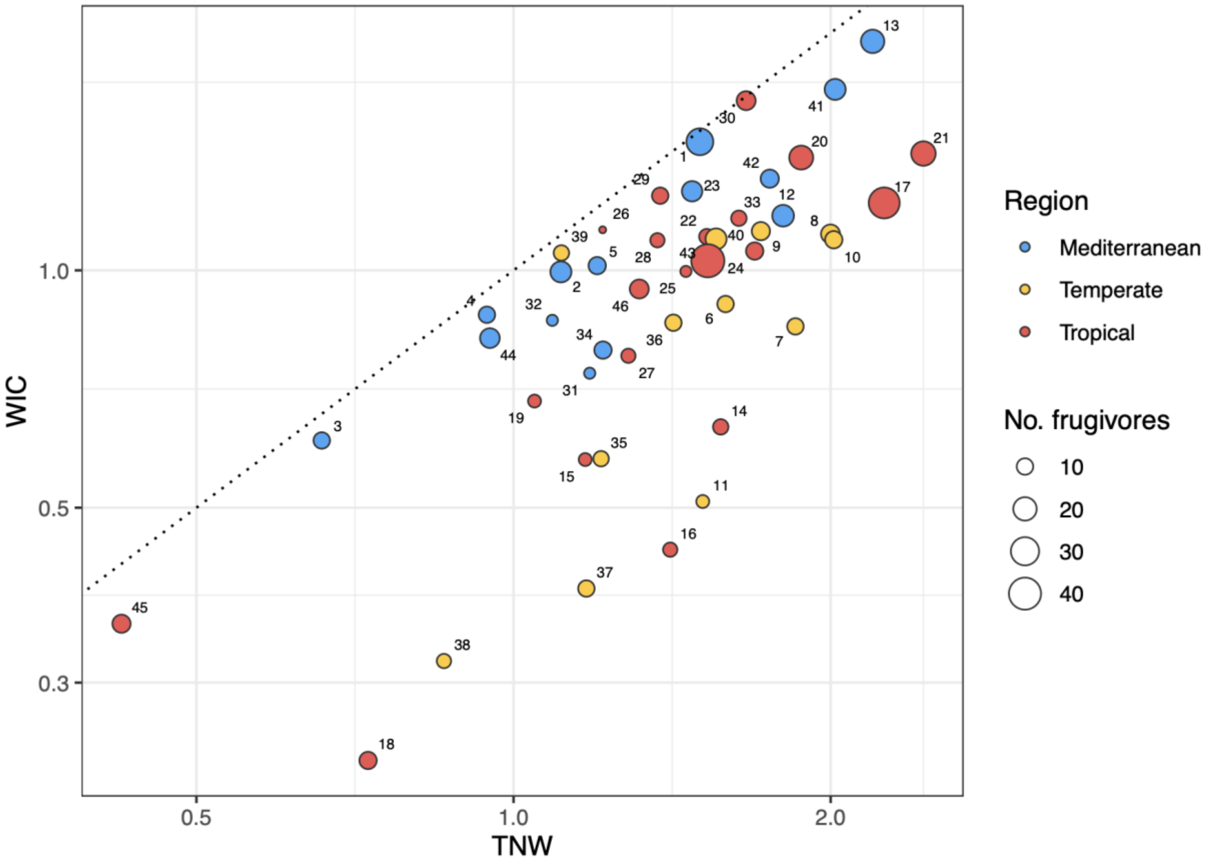


**Figure S5.** Within Individual Component (WIC) versus Total niche width (TNW) for individual-based frugivory networks. Point size is proportional to the number of observed frugivore species in the network, point color indicates the geographic region and number the network id (see Table S1). Note the log-scale in both axes. The dotted line represents a 1:1 ratio, in which the WIC would be equal to the TNW indicating individual niche widths that encompass the whole population niche width. The closer the networks are to the line, the higher WIC/TWN (i.e., lower individual specialization). Networks including many frugivore species tend to have a wider interaction niche (TNW), but not necessarily higher levels of individual specialization (WIC/TNW, i.e., far from the 1:1 line).

### Interaction curves by frugivores

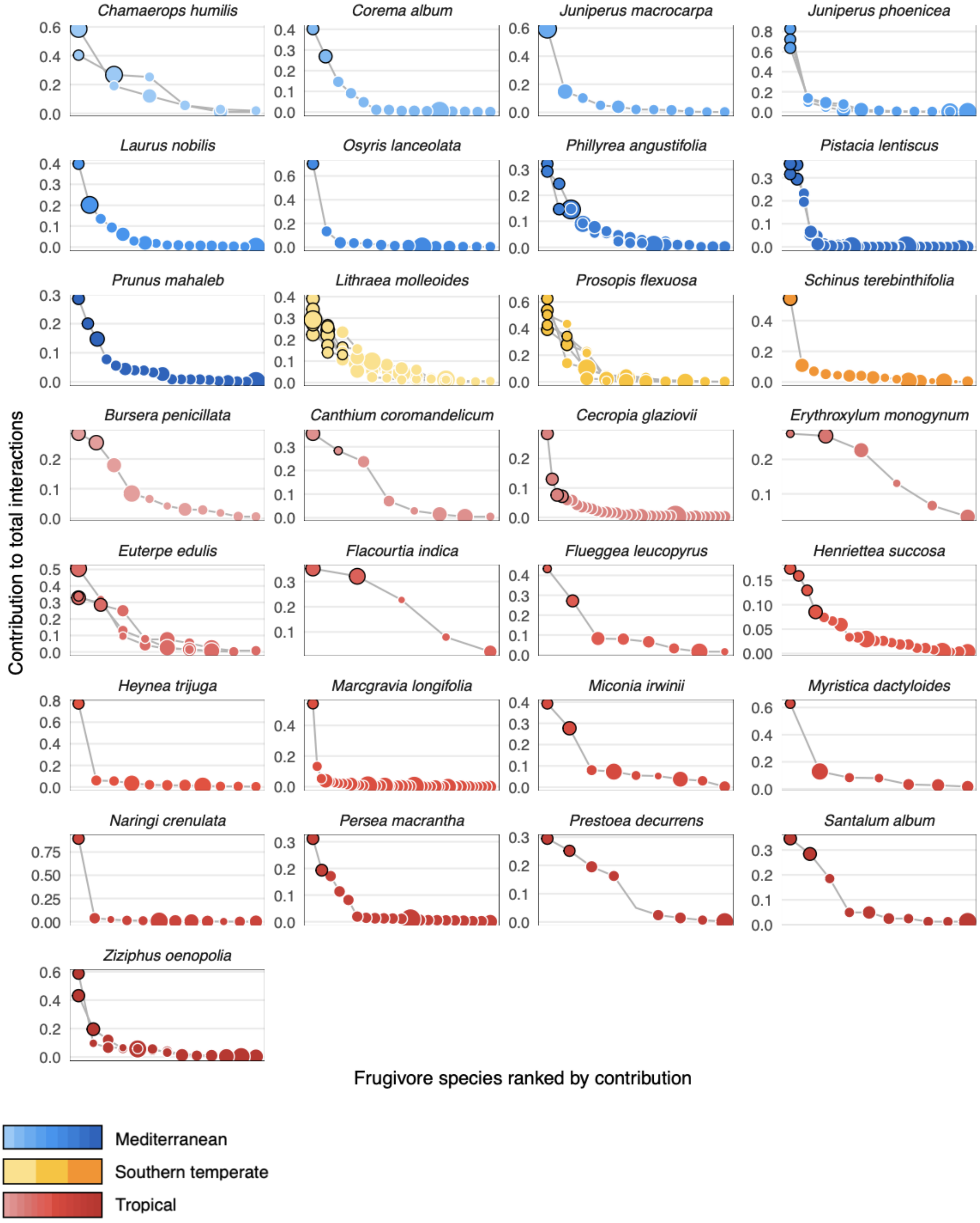


**Figure S6.** Relative contribution of each frugivore species (dots) to the total interactions of each plant species. Frugivores are ranked by decreasing contribution. Plant species with more than one population/network present several curves. Colors for each species correspond with different bioregions and different color shades differentiate plant species. Dots size represents frugivore body mass relative to the mean body mass of the assemblage (z-score) and black outlines in dots indicate those frugivore species whose aggregate contributions account for at least 50% of the interactions.

### Plant individuals' interaction profiles

#### *Node-level metrics*

Same as with network-level metrics, we tried to select node-level metrics that were not strongly correlated. All variables had a maximum VIF of 2.16 (VIF < 3).

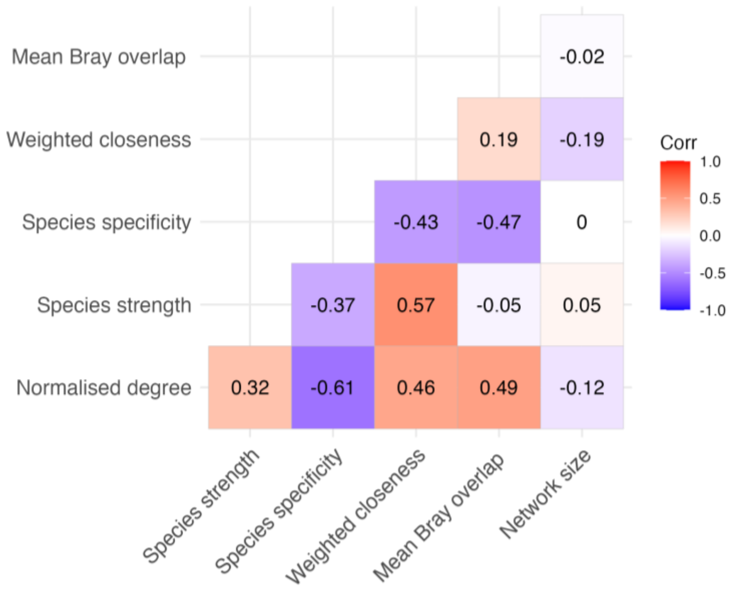


**Figure S7.** Correlation plot between selected node-level metrics. Numbers denote Pearson correlation coefficients (r) and the color its magnitude and direction.

*PCA analysis for comparing plant individuals' interaction profiles*

**Table S4.** Principal Component Analysis results node-level metrics.

| | PC1 | PC2 | PC3 | PC4 | PC5 |
|---|---|---|---|---|---|
| **Importance of components:** | | | | | |
| Eigenvalue | 1.60 | 1.11 | 0.71 | 0.59 | 0.58 |
| Proportion of Variance | 0.51 | 0.25 | 0.10 | 0.07 | 0.07 |
| Cumulative Proportion | 0.51 | 0.76 | 0.86 | 0.93 | 1.00 |
| **PC loadings:** | | | | | |
| Normalised degree | -0.52 | -0.20 | -0.23 | -0.72 | -0.34 |
| Species strength | -0.37 | 0.63 | -0.20 | 0.40 | -0.51 |
| Species specificity index | 0.51 | 0.15 | 0.5 | -0.28 | -0.61 |
| Weighted closeness | -0.45 | 0.38 | 0.7 | -0.17 | 0.36 |
| Mean Bray-Curtis overlap | -0.35 | -0.63 | 0.40 | 0.46 | -0.33 |

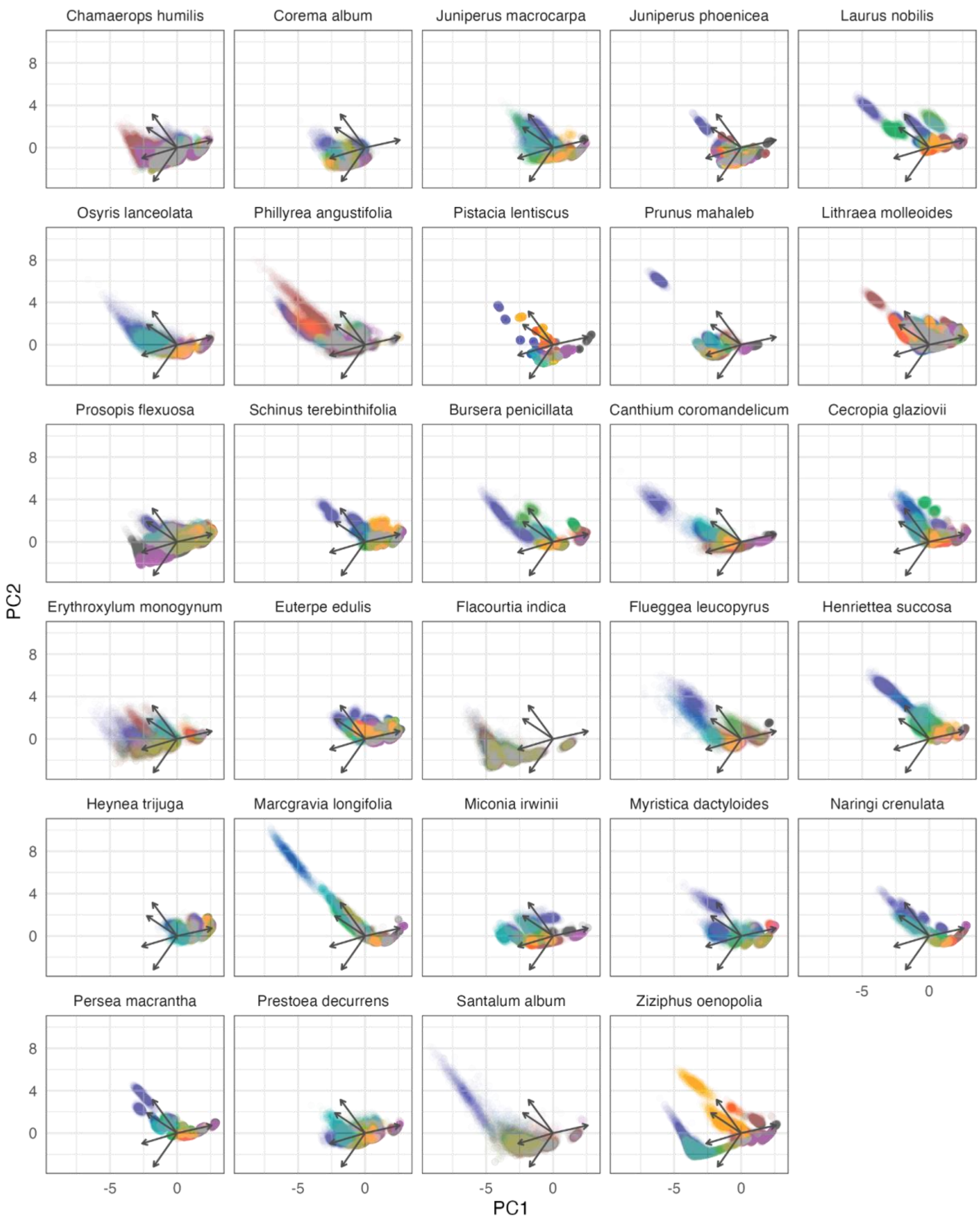


**Figure S8.** Principal Component Analysis for node-level metrics of individual plants in their respective networks. PCA multivariate space is faceted by plant species to facilitate display of plant individuals distribution in the multivariate space and the identification of outlying individuals. Note that some species present more than one population (i.e., more than one network, see Table S1). Each individual plant is represented by a point cloud with a different color and its node-level metrics were estimated using the full network posterior distribution (n = 1000 points per individual). It is visible how some individuals have well delimited point clouds, while some other individuals have much more uncertainty due to lower sampling coverage. See Fig. 5 for information on what node-level metric represents each of the five arrows.

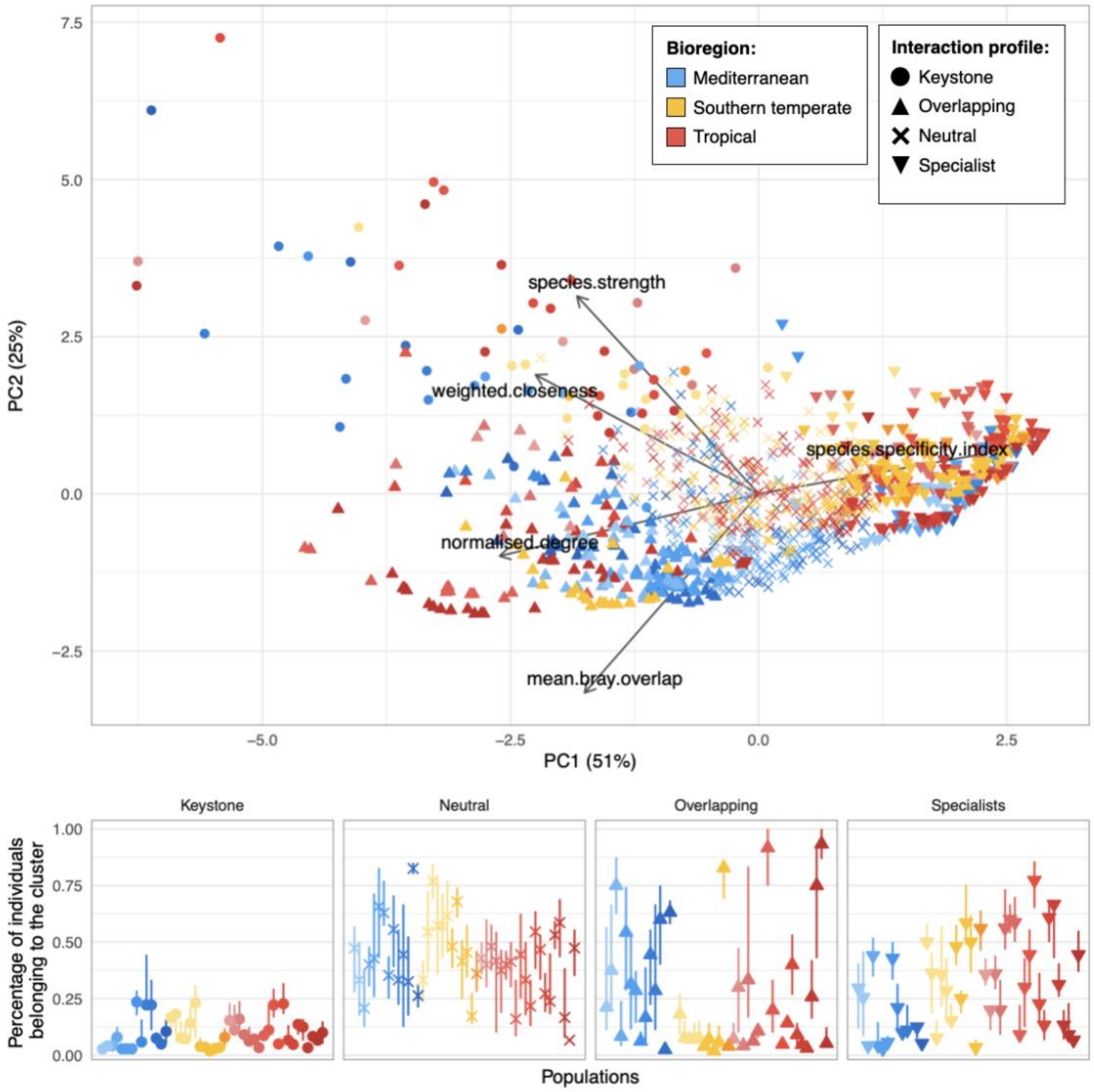


**Figure S9.** PCA for node-level metrics in all plant populations. Each point represents an individual plant. For each plant individual we have represented the centroid of its posterior distribution node-metric values, see Fig. S8. The shape of the points represent the most common categorization into one of the four different interaction profiles of each plant individual. The panel below shows the proportion of individuals within each population that fall into one of these clusters or interaction profiles. The interval of each point represents the 90% confidence interval for the 1000 cluster analysis repetitions. Colors indicate the bioregion and different shades refer to different plant species within each bioregion.

### Software citations

We used R version 4.4.0 (R Core Team 2024) and the following R packages: adegenet v. 2.1.10 (Jombart 2008; Jombart and Ahmed 2011), BayesianWebs v. 0.0.7 (Rodriguez-Sanchez 2024a), bayestestR v. 0.13.2 (Makowski, Ben-Shachar, and Lüdecke 2019), bipartite v. 2.19 (Dormann, Gruber, and Fruend 2008; Dormann et al. 2009; Dormann 2011), cluster v. 2.1.6 (Maechler et al. 2023), DHARMa v. 0.4.6 (Hartig 2022), fmsb v. 0.7.6 (Nakazawa 2024), GGally v. 2.2.1 (Schloerke et al. 2024), ggcorrplot v. 0.1.4.1 (Kassambara 2023), ggdist v. 3.3.2 (Kay 2024b, 2024a), ggfortify v. 0.4.17 (Tang, Horikoshi, and Li 2016; Horikoshi and Tang 2018), ggh4x v. 0.2.8 (van den Brand 2024), ggrepel v. 0.9.5 (Slowikowski 2024), ggridges v. 0.5.6 (Wilke 2024), glmmTMB v. 1.1.9 (Brooks et al. 2017), gridExtra v. 2.3 (Auguie 2017), here v. 1.0.1 (Müller 2020), igraph v. 2.0.3 (Csardi and Nepusz 2006; Csárdi et al. 2024), iNEXT v. 3.0.1 (Chao et al. 2014; Hsieh, Ma, and Chao 2024), knitr v. 1.46 (Xie 2014, 2015, 2024), MetBrewer v. 0.2.0 (Mills 2022), modelbased v. 0.8.7 (Makowski et al. 2020), network.tools v. 0.0.4 (Rodriguez-Sanchez 2024b), paletteer v. 1.6.0 (Hvitfeldt 2021), patchwork v. 1.2.0 (Pedersen 2024), psych v. 2.4.3 (William Revelle 2024), rcartocolor v. 2.1.1 (Nowosad 2018), renv v. 1.0.7 (Ushey and Wickham 2024), reshape2 v. 1.4.4 (Wickham 2007), rmarkdown v. 2.26 (Xie, Allaire, and Grolemund 2018; Xie, Dervieux, and Riederer 2020; Allaire et al. 2024), scales v. 1.3.0 (Wickham, Pedersen, and Seidel 2023), summarytools v. 1.0.1 (Comtois 2022), tidylog v. 1.0.2 (Elbers 2020), tidyverse v. 2.0.0 (Wickham et al. 2019), tnet v. 3.0.16 (Opsahl 2009).

Xie, Yihui. 2014. "knitr: A Comprehensive Tool for Reproducible Research in R." In Implementing Reproducible Computational Research, edited by Victoria Stodden, Friedrich Leisch, and Roger D. Peng. Chapman; Hall/CRC.

———. 2015. Dynamic Documents with R and Knitr. 2nd ed. Boca Raton, Florida: Chapman; Hall/CRC. https://yihui.org/knitr/.

———. 2024. knitr: A General-Purpose Package for Dynamic Report Generation in r. https://yihui.org/knitr/.

Xie, Yihui, J. J. Allaire, and Garrett Grolemund. 2018. R Markdown: The Definitive Guide. Boca Raton, Florida: Chapman; Hall/CRC. https://bookdown.org/yihui/rmarkdown.

Xie, Yihui, Christophe Dervieux, and Emily Riederer. 2020. R Markdown Cookbook. Boca Raton, Florida: Chapman; Hall/CRC. https://bookdown.org/yihui/rmarkdown-cookbook
